# The small and large intestine contain transcriptionally related mesenchymal stromal cell subsets that derive from embryonic *Gli1^+^* mesothelial cells

**DOI:** 10.1101/2021.08.13.456086

**Authors:** Simone Isling Pærregaard, Sophie Schussek, Line Wulff, Kristoffer Niss, Urs Mörbe, Johan Jendholm, Kerstin Wendland, Anna T. Andrusaite, Kevin F. Brulois, Robert J. B. Nibbs, Katarzyna Sitnik, Allan McI Mowat, Eugene C. Butcher, Søren Brunak, William W. Agace

**Author notes:** Equal contribution (Sophie Schussek, Line Wulff). Correspondence: William Agace.

## Abstract

Intestinal fibroblasts (FB) play essential roles in intestinal homeostasis. Here we show that the small and large intestinal lamina propria (LP) contain similar FB subsets that locate in specific anatomical niches and express distinct arrays of epithelial support genes. However, there were tissue specific differences in the transcriptional profile of intestinal FB subsets in the two sites. All adult intestinal LP mesenchymal stromal cells (MSC), including FB, smooth muscle cells (SMC) and pericytes derive from *Gli1*-expressing embryonic precursors which we identify as mesothelial cells. Trajectory analysis suggested that adult SMC and FB derive from distinct embryonic intermediates, and that adult FB subsets develop in a linear trajectory from CD81^+^ FB. Finally, we show that colonic subepithelial PDGFRα^hi^ FB comprise several functionally and anatomically distinct populations that originate from an *Fgfr2*-expressing FB intermediate. Collectively our results provide novel insights into MSC diversity, location, function and ontogeny, with implications for our understanding of intestinal development, homeostasis and disease.

## Introduction

The small and large intestines form a continuous tube from the stomach to the anus, but are functionally and anatomically distinct. The small intestine is the primary site of food digestion and nutrient absorption and is characterized by finger-like projections termed villi that protrude into the intestinal lumen and maximize the absorptive area of the epithelium. In contrast, the large intestine is primarily a site of water absorption and is a major niche for beneficial microbes; its surface consists of crypts linked by short regions of flat surface epithelium. The cellular composition of the intestinal mucosa also differs markedly between the small and large intestines (Mowat and Agace, 2014; Agace and McCoy, 2017). For example, the small and large intestines contain different numbers and proportions of innate and adaptive immune cells as well as epithelial subpopulations (Barker, 2014; Mowat and Agace, 2014; Agace and McCoy, 2017; Parikh *et al*., 2019). These distinct segments are also exposed to different concentrations of microbial and food-derived metabolites that regulate the composition and function of local cells (Mowat and Agace, 2014; Agace and McCoy, 2017). However, the cellular and signaling components that determine the differences in tissue structure and composition are not fully understood.

The intestinal lamina propria (LP) contains a large population of tissue resident mesenchymal stromal cells (MSC) that include fibroblasts (FB), pericytes (PC) and smooth muscle cells (SMCs) that play an essential role in intestinal homeostasis (Degirmenci *et al*., 2018; Kinchen *et al*., 2018; Shoshkes-Carmel *et al*., 2018; Brügger *et al*., 2020; Hong *et al*., 2020; McCarthy *et al*., 2020; Wu *et al*., 2021). For example, intestinal FB are major producers of extracellular matrix proteins that help provide structure to the mucosa (Furuya and Furuya, 2007; Roulis and Flavell, 2016). They also express factors essential for epithelial (Stzepourginski *et al*., 2017; Degirmenci *et al*., 2018; Shoshkes-Carmel *et al*., 2018; Karpus *et al*., 2019; McCarthy *et al*., 2020; Wu *et al*., 2021) and endothelial homeostasis (Thomson *et al*., 2018; Hong *et al*., 2020; Fawkner-Corbett *et al*., 2021), as well as immune cell localization and function (Fagarasan *et al*., 2001; Powell *et al*., 2011; Beswick *et al*., 2014; Vicente-Suarez *et al*., 2015). Recent single cell (sc)RNA-seq studies have demonstrated considerable heterogeneity within the intestinal LP MSC compartment and have led to the identification of several FB clusters with non-redundant functions in intestinal homeostasis (Kinchen *et al*., 2018; Smillie *et al*., 2019; Brügger *et al*., 2020; Hong *et al*., 2020; McCarthy *et al*., 2020; Roulis *et al*., 2020; Wu *et al*., 2021). A picture is also emerging whereby different intestinal FB subsets locate within distinct regions of the mucosa (Degirmenci *et al*., 2018; Shoshkes-Carmel *et al*., 2018; Thomson *et al*., 2018; Parikh *et al*., 2019; Hong *et al*., 2020; McCarthy *et al*., 2020; Fawkner-Corbett *et al*., 2021), providing specialized support to cells in their local environment (Aoki *et al*., 2016; Stzepourginski *et al*., 2017; Degirmenci *et al*., 2018; Kinchen *et al*., 2018; Shoshkes-Carmel *et al*., 2018; Karpus *et al*., 2019; Hong *et al*., 2020; McCarthy *et al*., 2020; Wu *et al*., 2021). However, the exact nature of these diverse LP MSC subsets and how they differ in the small and large intestine remains to be established.

scRNA-seq analyses have shown that the composition of human intestinal MSC populations changes markedly as the tissue develops in the embryo (Fawkner-Corbett *et al*., 2021; Holloway *et al*., 2021). Although the origin of these populations remains to be determined, lineage-tracing experiments in mice have suggested that the mesothelium, an epithelial monolayer that lines the serosal surface of the intestine (Koopmans and Rinkevich, 2018), can give rise to SMC and various FB in the intestinal serosa and muscle layers (Wilm *et al*., 2005; Rinkevich *et al*., 2012). Whether MSC subsets present within the adult intestinal LP derive from cells of common or distinct embryonic origin and the developmental relationship between adult MSC subsets remains unclear.

Here we demonstrate that LP MSC subset composition is similar in the small and large intestine and that each subset occupies distinct anatomical niches. Nevertheless, the transcriptional profile of the major LP FB subsets differed markedly between the small and large intestine, suggesting regional specific functions in intestinal homeostasis. Grafting and lineage-tracing experiments demonstrated that all MSC subsets in adult small intestinal and colonic LP derive from *Gli1*-expressing precursors present in embryonic day (E)12.5 intestine. Computational analysis suggested that all adult intestinal MSC derive from embryonic intestinal mesothelial cells and that adult SMC and FB arise from distinct mesothelial derived embryonic intermediates. We could also define a linear developmental trajectory for all adult FB subsets that originated from CD81^+^ FB.

## Results

### The small intestine and colon LP contain diverse, but transcriptionally related MSC subsets

To gain a broad understanding of MSC subset diversity in the intestinal LP, we performed scRNA-seq on MSC isolated from the small intestine and colon LP of 8-10 week old mice. Briefly, after removal of Peyer’s patches, muscularis externa and epithelium, intestinal MSCs were enriched from digested intestinal LP cell suspensions by fluorescently activated cell sorting of live, single, lineage^-^ (CD45^+^, Ter119^+^), non-epithelial (EpCAM^+^), non-endothelial (CD31^+^), non-lymphoid tissue-associated MSCs (BP3^+^) (Taylor *et al*., 2007) and non-glial cells (L1CAM^+^), followed by gating on cells expressing the pan MSC marker Itgβ1 (Fig. S1A). After bioinformatic removal of contaminating c-kit^+^ interstitial cells of Cajal (ICC), CD31^+^ endothelial cells, plasma cells and CD45^+^ immune cells, sequencing data of 16.964 small intestinal and 14.164 colonic MSC remained.

Louvain clustering identified six small intestinal MSC clusters (Fig. 1A) and differential gene expression (DEG) analysis of these clusters identified pericytes, SMC and four FB clusters (Fig. S1B). These were PDGFRα^hi^ FB, two PDGFRα^lo^CD34^hi^ clusters that could be distinguished based on their expression of *Cd81* (hereafter called CD81^+^ FB) and *Igbp5* (hereafter called Igfbp5^+^ FB), and a PDGFRα^lo^CD34^lo^ cluster that expressed higher levels of *Fgfr2* (hereafter called Fgfr2^+^ FB) (Fig. S1C). To determine how these clusters might relate to those identified in other, recently published scRNA-seq studies of small intestinal MSC (Hong *et al*., 2020; McCarthy *et al*., 2020), DEGs from the previous MSC subsets were overlaid with our scRNA-seq dataset (Fig. S1D). The MSC population termed “mural cells” by Hong *et al* (Hong *et al*., 2020) corresponded to our pericytes, while their FB subsets termed FB2, 3, 4 and 5 corresponded to our small intestinal Igfbp5^+^ FB, Fgfr2^+^ FB, CD81^+^ FB and PDGFRα^hi^ FB clusters, respectively (Fig. S1D). The signature genes of FB1 identified by Hong *et al* as activated FB based on their expression of *Junb* and *Fosb*, were expressed widely by several MSC subsets in our dataset (Fig. S1D), indicating that this cluster represents a cell state rather than an MSC subset. Similar analysis of the MSC datasets generated by McCarthy *et al* (McCarthy *et al*., 2020) demonstrated that the PDGFRα^hi^ MSC subset they defined as “telocytes” corresponded to our PDGFRα^hi^ FB cluster, while their Lo-1 FB subset corresponded to our CD81^+^ FB cluster and their Lo-2 FB subset encompassed both our Fgfr2^+^ and Igfbp5^+^ FB clusters (Fig. S1D) (McCarthy *et al*., 2020). Thus, our results confirm and extend recent findings and highlight the complexity of MSC subsets in the small intestinal LP. Louvain clustering also identified six MSC clusters in colon LP (Fig. 1B), which DEG analysis identified as pericytes, SMC, and four FB clusters (Fig. S1E). These were PDGFRα^hi^ FB and three PDGFRα^lo^CD34^+^ clusters that could be distinguished based on their expression of *Cd81* (hereafter called CD81^+^ FB), *CD90* (hereafter called CD90^+^ FB) or *Fgfr2* (hereafter called Fgfr2^+^ FB) (Fig. S1F). To determine the relationship between the colonic and small intestinal MSC subsets, Pearson correlation analysis was performed based on the pseudo-bulk of overlapping variable genes between the two data sets. This showed that colonic pericytes, SMC, PDGFRα^hi^ FB, CD81^+^ FB closely correlate with their counterparts in the small intestine, that colonic CD90^+^ FB most closely correlate with small intestinal Igfbp5^+^ FB and that colonic Fgfr2^+^ FB closely correlate with both Fgfr2^+^ and Igfbp5^+^ FB (Fig. 1C).

**Figure 1.**
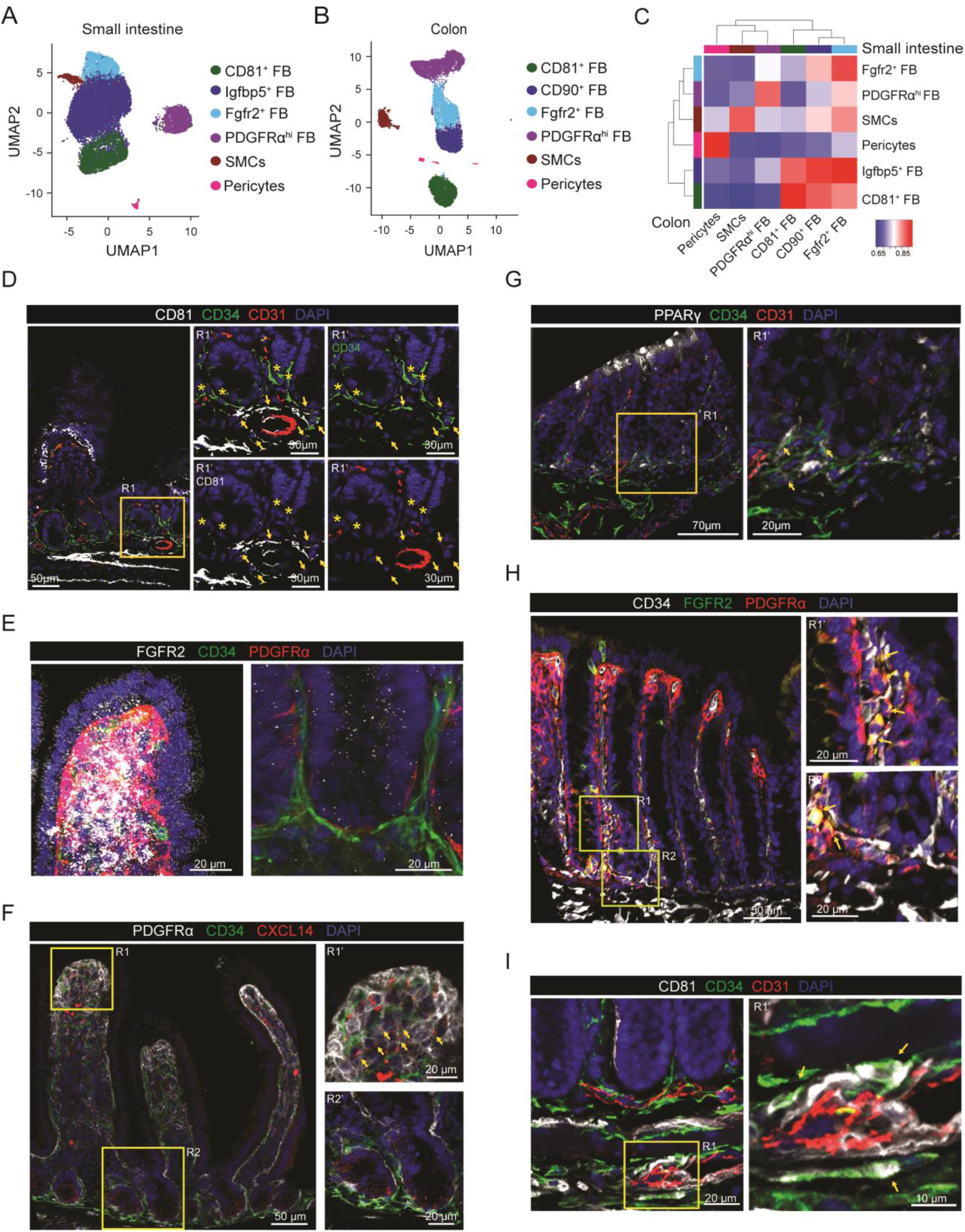
Intestinal MSC subsets are broadly conserved across intestinal segments. (**A-B**) Uniform Manifold Approximation and Projection (UMAP) colored by unsupervised Louvain clustering of murine small intestinal (**A**) and (**B**) colonic MSC. Results are from 2 independent experiments/organ with 3 pooled mice/experiment. (**C**) Pearson correlations between averaged cluster expressions of Louvain clusters from small intestinal and colonic MSC based on 1301 overlapping variable genes. Unsupervised hierarchical clustering indicate similarity of subsets within each tissue. (**D**-**I)** Immunohistochemical staining of mouse jejunum (**D-F**) or colon (**G-I**) for indicated antigens. (**D, F-I**) Region (R)1’ and R2’ represent magnifications of R1 and R2 quadrants (yellow squares). (**D**) Arrows indicate location of CD81^+^ FB (CD81^+^CD34^+^CD31^-^ cells) and stars, location of Igfbp5^+^ FB (CD81-CD34+ CD31^-^ cells). (**E**) Images of villus tip (left) and crypt (right). (**F-I**) Arrows indicate location of (**F**) Fgfr2^+^ FB (CXCL14^+^PDGFRα^+^CD34^-^ cells), (**G**) CD90^+^ FB (PPARγ^+^CD34^+^CD31^-^ cells), (**H**) Fgfr2^+^ FB (Fgfr2^+^CD34^+^PDGFRα^+^ cells) and (**I**) CD81^+^ FB (CD81^+^CD34^+^CD31^-^ cells). Results are representative stains from (**D-F, H**) 2 and (**G** and **I**) 3 experiments. See also Figure S1.

### FB subsets are located in distinct niches along the crypt-villus axis

There is increasing evidence that subsets of small intestinal FBs may occupy distinct anatomical niches that overlap with the WNT/BMP signaling gradient along the crypt-villus axis (Shoshkes-Carmel *et al*., 2018; Bahar Halpern *et al*., 2020; McCarthy *et al*., 2020). In line with a recent report (Stzepourginski *et al*., 2017), we found that small intestinal CD34^+^ FB (which include CD81^+^ and Igfbp5^+^ FB) were located around crypts and in the submucosa, but were largely excluded from the villus core (Fig. S1G). Of these, CD34^+^CD81^+^ FB located around CD31^+^ vessels close to and within the submucosa, with some locating close to crypts (Fig. 1D), consistent with recent reports (Thomson *et al*., 2018; Hong *et al*., 2020; McCarthy *et al*., 2020), while Igfbp5^+^ (CD34^+^CD81^-^) FB located around crypts (Fig. 1D). Conversely, PDGFRα^+^CD34^-^ FB (including both PDGFRα^hi^ FB and PDGFRα^lo^Fgfr2^+^ FB) were located directly underlying the epithelium and within the villus core (Fig. S1G). Of these, the Fgfr2^+^ FB were located within the villus core towards the tip of the villus (Fig. 1E); this was confirmed using CXCL14 as a marker for this subset (Fig. S1H and Fig. 1F). In contrast, the PDGFRα^hi^ FB lay directly under the epithelium and at the villus tip (Fig. 1E, Fig. 1F, Fig. S1G), supporting previous findings (Kurahashi *et al*., 2013; Bahar Halpern *et al*., 2020; Brügger *et al*., 2020; Hong *et al*., 2020; McCarthy *et al*., 2020).

As in the small intestine, colonic CD34^+^ FB subsets located beneath and surrounding intestinal crypts, while PDGFRα^hi^ FB formed a thin layer directly underlying the epithelium and were concentrated at the top of crypts (Fig. S1I). Colonic CD90^+^CD34^+^ FB, which expressed high levels of *Pparg* (Fig. S1H) could be identified after staining for PPARγ and were located at the base of colonic crypts (Fig. 1G). Colonic Fgfr2^+^ FB localized preferentially between crypts (Fig. 1H), whereas the colonic CD81^+^ FB located below the crypts and in the submucosa (Fig. 1I). Collectively these results demonstrate that the FB subsets identified by scRNA-seq locate within distinct niches of the small and large intestine.

### Expression of epithelial support genes is conserved across FB subsets in the small intestine and colon

Recent studies have suggested a division of labor amongst small intestinal FB subsets in the production of epithelial support factors (Degirmenci *et al*., 2018; Kinchen *et al*., 2018; Brügger *et al*., 2020; McCarthy *et al*., 2020; Wu *et al*., 2021) and we thus assessed the expression of such genes in our small intestinal and colonic FB datasets. Consistent with previous studies (Shoshkes-Carmel *et al*., 2018; McCarthy *et al*., 2020), small intestinal PDGFRα^hi^ FB were major producers of BMPs and this property was shared by colonic PDGFRα^hi^ FB (Fig. 2A). *Bmp3*, *Bmp5* and *Bmp7* expression was largely restricted to PDGFRα^hi^ FB, while expression of *Bmp1*, *Bmp2* and *Bmp4* was found more broadly among FB MSC subsets in both tissues (Fig. 2A). Both small intestinal and colonic PDGFRα^hi^ FB were also the dominant source of the non-canonical WNT ligands, *Wnt4*, *Wnt5a* and *Wnt5b*, although Fgfr2^+^ FB also expressed *Wnt4*, particularly in the small intestine (Fig. 2A). Consistent with previous results (Brügger *et al*., 2020; McCarthy *et al*., 2020), CD81^+^ FB were the major source of the BMP antagonist *Grem1* in the small intestine and this was also highly expressed by colonic CD81^+^ FB. However, in the colon, Fgfr2^+^ and CD90^+^ FB also expressed *Grem1* (Fig. 2A). Thus, the specialization of MSC subsets in their expression of epithelial support genes is largely conserved between the small intestine and colon.

**Figure 2.**
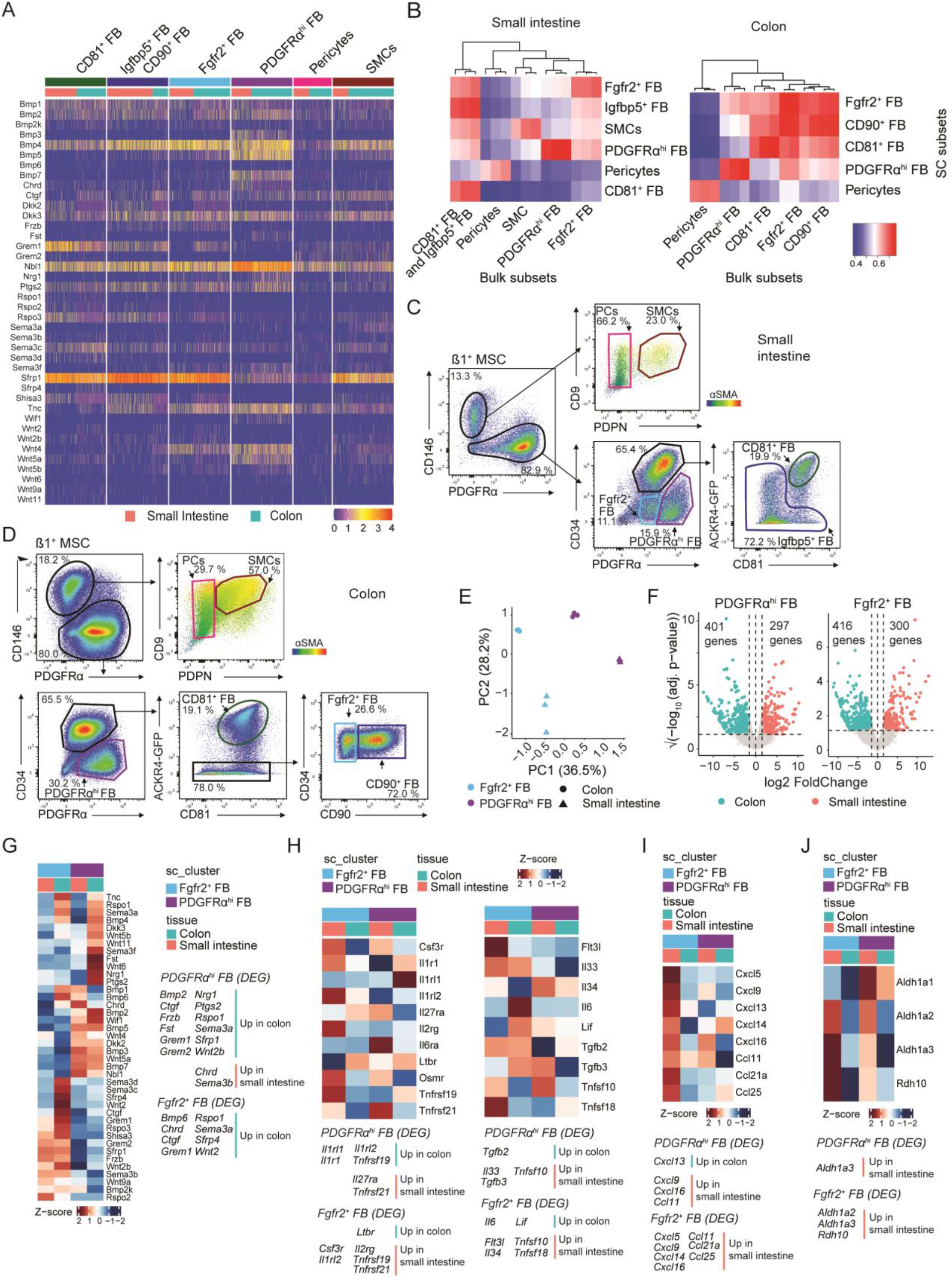
Despite similar FB subset composition, small intestinal and colonic FB display regional transcriptional specialization. (**A**) Heatmaps showing scaled expression (integrated data) of selected epithelial support genes by indicated MSC subsets. (**B**) Pearson correlations between averaged cluster expressions of Louvain clusters from scRNA-seq and bulk RNA-seq datasets based on 1937 (small intestine, left) and 1925 (colon, right) overlapping variable genes. Bulk RNA-seq data is from sorted MSC subsets from 3 independent experiments. Unsupervised hierarchical clustering indicate similarities of bulk RNA-seq subsets within each tissue. (**C-D**) Flow cytometric analysis of adult small intestinal (**C**) and colonic (**D**) Itgβ1^+^ MSCs from *Ackr4.GFP* mice. Representative staining of 2 experiments with 2-4 mice/experiment. Colored gates represent indicated MSC subsets. PCs - pericytes; SMCs - smooth muscle cells, FB - fibroblast. (**E**) Principal component analysis (PCA) of bulk RNA-seq data from indicated sorted FB populations. Results are from 3 independent sorts/population. (**F**) Volcano plots showing differentially expressed genes (DEGs) between small intestinal and colonic PDGFRα^hi^ FB (left) and Fgfr2^+^ FB (right). Dotted horizontal line denotes significant adjusted p-value of 0.05, vertical dotted lines denote log_2_FC = 0 and the log_2_FC of +/− 1.5. (**G-J**) Heatmap representations of averaged transcription levels of indicated genes within sorted FB subsets. Data are averaged from 3 independent bulk RNA-seq datasets. (**G**) Epithelial support genes, (**H**) cytokines and cytokine receptors, (**I**) chemokines, (**J**) vitamin A metabolism. Gene lists for **I** were selected based on the epithelial support list in (**A**) while those in **H-J** were differentially expressed between either small intestinal and colonic PDGFRα^hi^ FB or between small intestinal and colonic Fgfr2^+^ FB. Identified DEG that are 1.5<|log_2_FC| are listed to the right of (**G**) or below (**H-J**) the heat maps and. See also Figure S2.

### Small intestinal and colonic PDGFRα^hi^ FB and Fgfr2^+^ FB display regional transcriptional specificity

To gain a broader understanding of how the major PDGFRα^hi^ and Fgfr2^+^ FB subsets in the small and large intestine might be related, we examined our scRNA-seq datasets for surface markers that would allow us to identify and sort these cells for bulk RNA-seq analysis (Fig. S2A). For the small intestine, pericytes were identified and sorted as PDGFRα^-^ESAM- 1^+^PDPN^-^ cells, SMC as PDGFRα^-^ESAM-1^+^PDPN^+^ cells, PDGFRα^hi^ FB as ESAM-1^-^ PDGFRα^hi^CD34^-^ cells, Fgfr2^+^ FB as ESAM-1^-^PDGFRα^int^CD34^-^ cells and CD34^+^ FB (including CD81^+^ and Igfbp5^+^ FB) as ESAM-1^-^PDGFRα^int^CD34^+^ cells. For the colon, pericytes were sorted as for small intestine, PDGFRα^hi^ FB was sorted as ESAM-1^-^ PDPN^+^CD34^-^ cells, Fgfr2^+^ FB were sorted as ESAM-1^-^PDPN^hi^CD34^+^CD90^-^ cells, CD90^+^ FB as ESAM-1^-^CD34^+^PDPN^hi^CD90^+^ cells and CD81^+^ FB as ESAM-1^-^PDPN^int^CD34^+^CD90^-^ cells, based on the fact that colonic CD81^+^ FB express low levels of PDPN compared with the other CD34^+^ colonic FB subsets (Fig. S2B). Correlation analysis of these bulk sorted intestinal FB subsets with the scRNA-seq data confirmed the accuracy of this staining strategy to identify small intestinal and colonic FB subsets by flow cytometry (Fig. 2B and Fig S2C). This initial panel was then refined for use in subsequent flow cytometry based analysis by including anti- CD81 to positively identify CD81^+^ FB directly, together with anti-CD146 (Fig. 2C and D), which can be used interchangeably with ESAM-1 (Fig. S2D). CD81^+^ FB also expressed the atypical chemokine receptor, ACKR4, as assessed using *Ackr4*.GFP reporter mice (Fig. 2C and D) (Thomson *et al*., 2018), consistent with previous reports (Thomson *et al*., 2018; Brügger *et al*., 2020; Hong *et al*., 2020; McCarthy *et al*., 2020) and our scRNA-seq analysis (Fig. S1B).

PCA analysis of bulk sorted PDGFRα^hi^ FB and Fgfr2^+^ FB distinguished these subsets from one another in PC1, while PC2 separated small intestinal from colonic FB (Fig. 2E), suggesting that anatomical location has a major impact on the transcriptional profile of these FB subsets. Consistent with this, small intestinal and colonic PDGFRα^hi^ FB differed in their transcription of 698 genes, while the two populations of Fgfr2^+^ FB differed in their transcription of 716 genes (Fig. 2F, see Supplementary Table 1 (for PDGFRα^hi^ FB) and Supplementary Table 2 (for Fgfr2^+^ FB) for complete list). Of these, 149 genes were differentially expressed between the small intestine and colon in both FB subsets (Supplementary Table 3); this included numerous Hox genes (Fig. S2E), consistent with the role of mesoderm in specifying the development of the different intestinal segments (Yuasa, 2003). Enrichr based analysis (Bioplanet 2019 (Huang *et al*., 2019)) showed that most of the upregulated pathways in the two subsets were in colon compared with small intestine, with few being upregulated in small intestine compared with colon (Fig. S2F). Irrespective of their location, PDGFRα^hi^ FB and Fgfr2^+^ FB showed very distinct expression of epithelial support genes (Fig. 2G), suggesting these populations play discrete roles in maintaining the epithelium; many of these genes were expressed at significantly higher levels in colonic subsets compared with their small intestinal counterparts (Fig. 2G). PDGFRα^hi^ FB and Fgfr2^+^ FB also expressed a wide range of immunologically relevant genes in both a subset- and tissue-specific manner (Fig. 2H-J). This included several cytokine and cytokine receptors (Fig. 2H), while small intestinal but not colon Fgfr2^+^ FB expressed a wide range of chemokines (Fig. 2I). Both subsets of small intestinal FB also expressed enzymes implicated in vitamin A metabolism, including the generation of retinoic acid (Fig. 2J), a major regulator of small intestinal immune responses. Collectively, these results highlight the unique functions of intestinal PDGFRα^hi^ FB and Fgfr2^+^ FB and show that these vary depending on anatomical location.

### Intestinal precursors in E12.5 intestine can give rise to all adult intestinal MSC subsets

While adult small intestinal and colonic LP contains multiple phenotypically, transcriptionally, and spatially distinct MSC subsets, the developmental relationship between these subsets and whether all derive from similar precursors remains unclear (Wilm *et al*., 2005; Rinkevich *et al*., 2012; Fawkner-Corbett *et al*., 2021). To explore this, we first investigated which MSC might be present in the small intestine and colon of E12.5 embryos by flow cytometry (Fig. 3A and S3A). In contrast to adult mice (Fig. 2C and D), E12.5 small intestinal and colonic Itgβ1^+^ cells consisted of one major population of PDGFRα^+^CD34^-^ MSCs, together with a small subset of PDGFRα^-^ cells that expressed the mesothelial markers dipeptidyl peptidase-4 (DPP4, CD26) and PDPN (Fig. 3A and B) (Meyerholz, Lambertz and McCray, 2016). To assess whether these populations could give rise to the MSC subsets found in the adult intestine, small and large intestine were dissected from E12.5 embryos and transplanted under the kidney capsule of adult WT recipient mice (Fig. 3C). Embryonic intestines from mice ubiquitously expressing EYFP were used for these experiments in order to trace the development of donor derived (EYFP^+^) MSC within grafted tissues. As expected (Ferguson, Parrott and Connor, 1972; Mosley and Klein, 1992; Yanai *et al*., 2017), small intestinal and colonic grafts had increased markedly in size by 4-6 weeks post transplantation (Fig. 3C) and contained mucosa that histologically resembled that of adult small intestine and colon, respectively (Fig. S3B). To assess the phenotypic diversity of graft-derived MSC, small intestinal and colonic grafts were isolated 4 weeks after transplantation, digested, and the expression of MSC subset markers on embryonically derived (YFP^+^) Itgβ1^+^ MSC assessed by flow cytometry (Fig. 3D, Fig S3C). Both small intestinal and colonic grafts contained putative populations of graft-derived SMC (CD146^+^PDGFRα^-^PDPN^+^), pericytes (CD146^+^PDGFRα^-^PDPN^-^), PDGFRα^hi^ FB (CD146^-^ CD34^-/lo^PDGFRα^hi^), CD81^+^ FB (CD146^-^PDGFRα^lo^CD81^+^CD34^+^) and CD81^-^CD34^+^ FB (CD146^-^PDGFRα^lo^CD34^+^CD81^-^) (Fig. 3D). To confirm the presence of these MSC subsets in the grafts, YFP^+^Itgβ1^+^ MSC were sorted from grafted colon and subjected to scRNA-seq (Fig. S3C). UMAP dimensionality reduction and Louvain clustering identified eight clusters (Fig. 3E), two of which (clusters 6 and 7) were identified as ICC and mesothelial cells, respectively (Kanamori-Katayama *et al*., 2011; Lee *et al*., 2017; Namvar *et al*., 2018) (Fig. S3D). These clusters were not part of our adult MSC datasets, as ICC were removed bioinformatically and the mesothelium was removed together with the muscularis externa during tissue processing. Pearson correlation analysis based on the pseudobulk of overlapping variable genes identified cluster 3 as being similar to adult PDGFRα^hi^ FB, cluster 4 as pericytes and cluster 5 as SMC (Fig. 3E and F). The remaining three clusters (clusters 0-2) were more closely related to the three adult CD34^+^ FB subsets, with cluster 1 most closely correlated to CD81^+^ FB, cluster 0 most closely correlated to Fgfr2^+^/CD90^+^ FB and cluster 2 showing equivalent correlation to all three adult CD34^+^ FB subsets (Fig. 3F). Furthermore, the distinct expression of epithelial support genes by each of the four FB subsets largely overlapped with the pattern seen in adult intestine (Fig. 3G, Fig. 2A). Collectively, these results suggest that embryonal MSC precursors present in E12.5 intestine can give rise to all adult intestinal MSC subsets.

**Figure 3.**
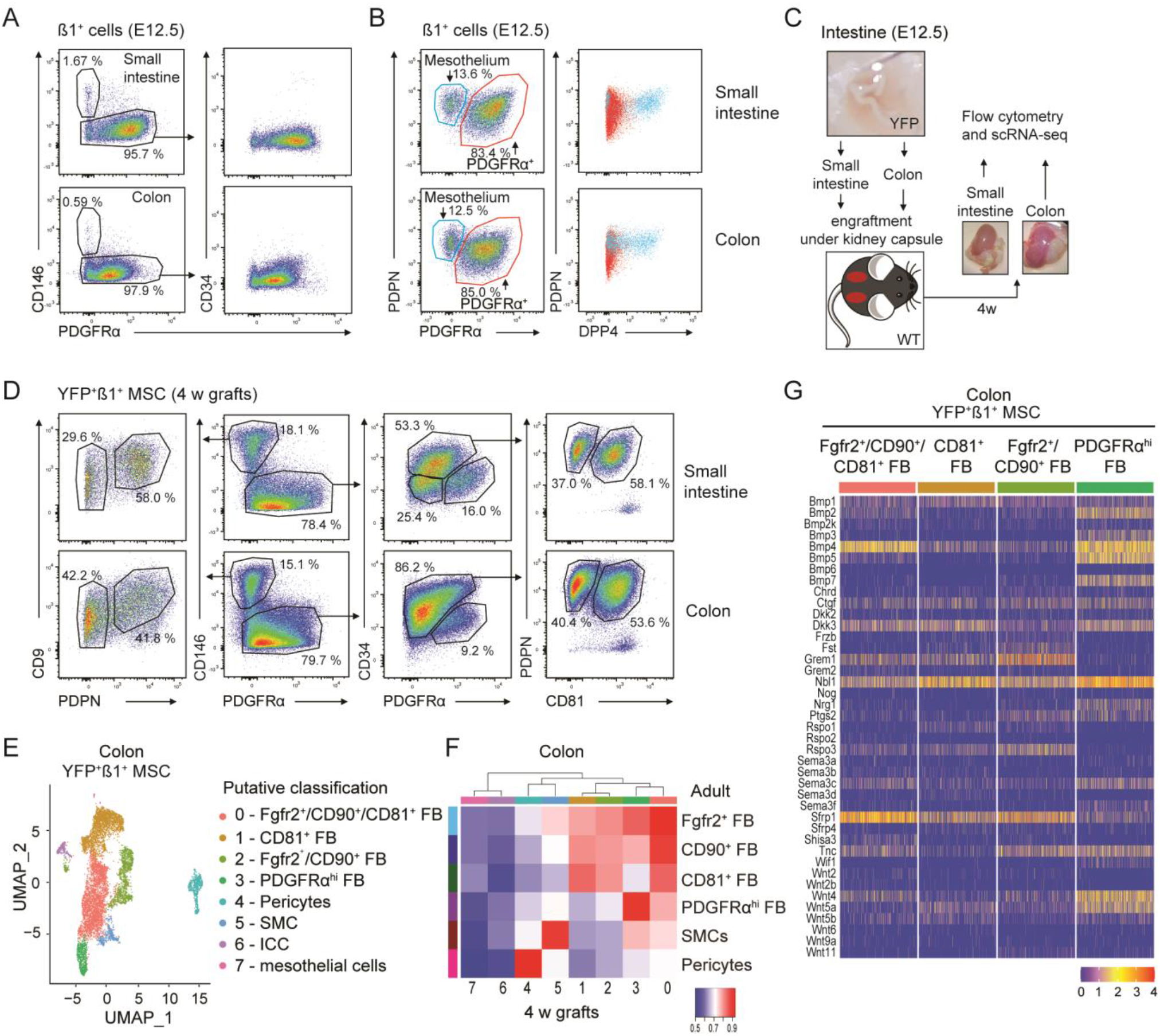
Adult intestinal MSC subsets derive from intestinal precursors present in E12.5 intestine. (**A-B**) Flow cytometric analysis of Itgβ1^+^ MSCs isolated from indicated organs on embryonic day (E) 12.5. (**B**) Right hand plots show expression of DPP4 (CD26) on gated PDPN^+^PDGFRα^-^ (blue) and PDPN^+^PDGFRα^+^ (red) cells from plots on left. Data are representative of (**A**) 4 experiments with 2-8 embryos/experiment, or (**B**) 3 experiments with 6-8 individual embryos. (**C**) Workflow of transplantation of E12.5 intestine from YFP^+^ mice under the kidney capsule of WT recipients. (**D**) Flow cytometric analysis of YFP^+^Itgβ1^+^ MSC in intestinal grafts 4 weeks after transplantation. Results are representative of 2 experiments with 4 (small intestine) or 2-3 (colon) grafts/experiment. (**E**) UMAP dimensionality reduction of scRNA-seq data colored by Louvain clustering from FACS purified YFP^+^Itgβ1^+^ MSC isolated from colonic grafts 4 weeks after transplantation. Data are from 8624 single cells from 3 pooled colonic grafts with an average of 2223 genes/cell. (**F**) Pearson correlations of averaged gene expression in colonic graft and adult colon MSC clusters based on 1486 overlapping variable genes. (**G**) Heatmap showing scaled transcription levels (integrated data) of selected epithelial support genes within the putative corresponding FB clusters identified in (**E**). See also Figure S3.

### Adult intestinal MSC derive from *Gli1^+^* embryonic precursors

To explore further the origin of adult MSC, we next lineage-traced E12.5 MSC and mesothelium into adulthood. GLI1 is a transcription factor induced by active hedgehog-signaling and is expressed by MSC in multiple organs (Kramann *et al*., 2015). PDGFRα^+^CD34^-^ MSC and PDGFRα^-^PDPN^+^ mesothelial cells from the small intestine and colon of E12.5 *Gli1*- EGFP embryos both expressed EGFP, whereas intestinal epithelial, endothelial and CD45^+^ cells did not (Fig. 4A and B). To lineage trace *Gli1*-expressing cells into adulthood, female *R26R*.EYFP mice (Srinivas *et al*., 2001) were mated with *Gli1*-Cre.ERT2 males expressing the estrogen receptor (ERT2) under control of *Gli1*-Cre, and pregnant dams injected i.p with 4- hydroxytamoxifen (4-OHT) at E11.5 (Fig. 4C). Two days later, YFP expression had been induced in a small but consistent proportion of Itgβ1^+^PDGFRα^+^ MSC and mesothelial cells in the small intestine and colon of *Gli1*-CreERT2^+/-^.*R26R*.EYFP embryos, but not in Cre^-^ embryos (Fig. S4A and B). Labeling was not observed in intestinal epithelial, endothelial, or CD45^+^ immune cells of *Gli1*-CreERT2^+/-^.*R26R*.EYFP mice and thus was specific to intestinal MSC and mesothelial cells (Fig. S4A and B). 5-7 weeks after birth, similar proportions of YFP- expressing cells were detected in all mature MSC subsets in both the small intestine and colon (Fig. 4D). Collectively, these results demonstrate that *Gli1^+^* cells present in the E12.5 intestine contain cells that give rise to all major adult intestinal MSC subsets.

**Figure 4.**
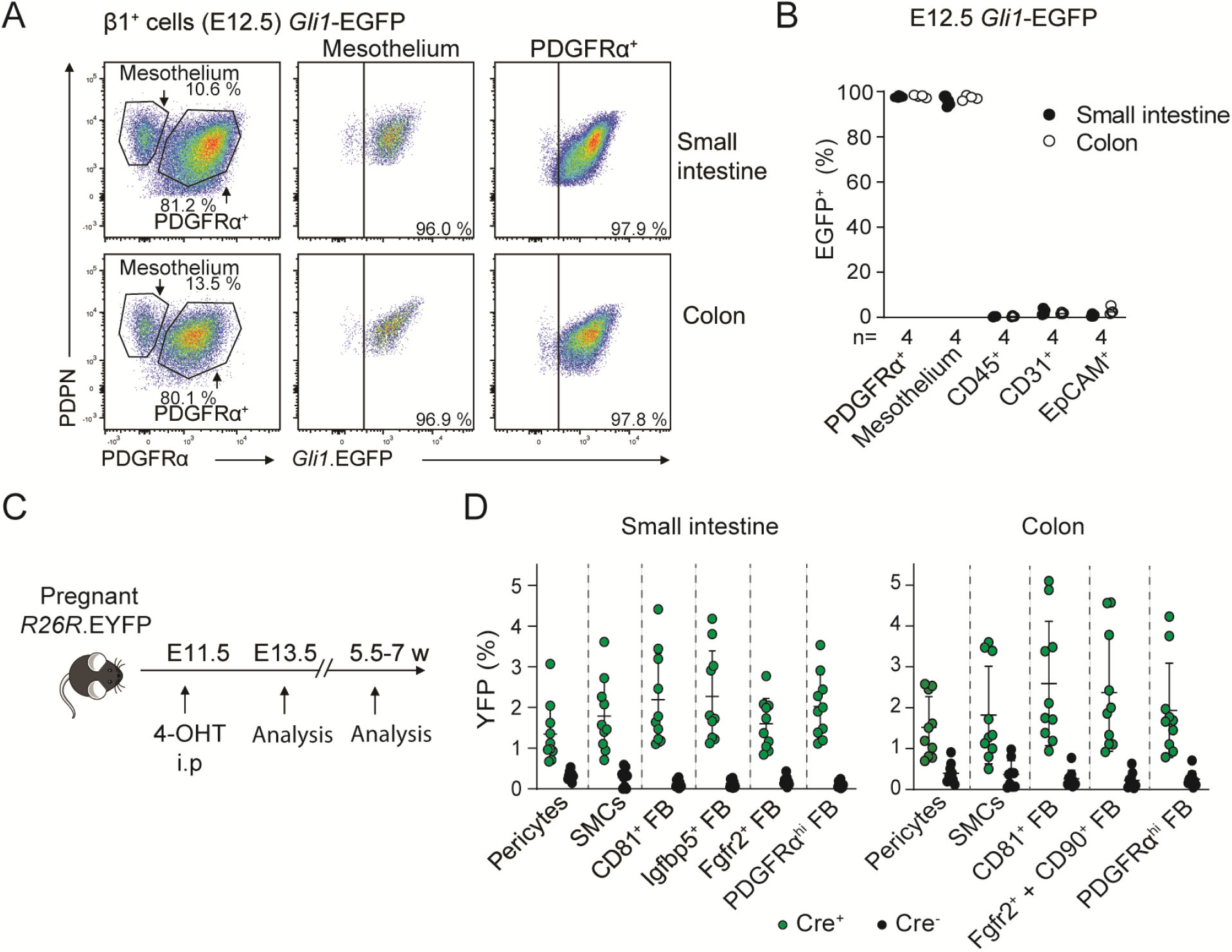
Adult intestinal MSC derive from *Gli1^+^* embryonic precursors. (**A**) Representative flow cytometric analysis and (**B**) Proportions of indicated cells expressing EGFP in the small intestine and colon of embryonic E12.5 *Gli1*-EGFP mice. Results are from 8 individual embryos, with each circle representing an individual embryo. (**C**) Workflow of lineage-tracing experiments. *R26R*.EYFP females were mated overnight with *Gli1*.CreERT2^+/-^ males and pregnant dams injected i.p. with 4-Hydroxytamoxifen (4-OHT) at E11.5. (**D**) Proportions of indicated MSC subset expressing YFP in small intestine and colon of 5.5-7 week old *Gli1*.CreERT2^+/-^.*R26R*.EYFP and *Gli1*.CreERT2^-/-^*R26R*.EYFP littermates. Results are from 4 independent experiments with 2-8 mice/experiment. Each circle represents an individual mouse. Bars represent the means and SD. See also Figure S4.

### Trajectory analysis indicates that adult intestinal MSC subsets originate from embryonic *Gli1^+^* mesothelial cells

To gain further insights into the relationship between embryonic intestinal *Gli1^+^* cells and adult intestinal MSC subsets, scRNA-seq was performed on fluorescently activated cell sorted Itgβ1^+^ MSC from the colon of E12.5 embryos. Louvain clustering identified six clusters (Fig. 5A), one of which, cluster 4, was identified as mesothelial cells due to its expression of mesothelial associated markers (Kanamori-Katayama *et al*., 2011; Namvar *et al*., 2018) (Fig. 5B). Consistent with our flow cytometric analysis (Fig. 3B), this cluster expressed transcripts for DPP4 and PDPN, but lacked expression of PDGFRα (Fig. S5A). To determine the relationship between embryonic and adult MSC subsets, the embryonic and adult colonic datasets were integrated and tSPACE (Dermadi *et al*., 2020) trajectory analysis was performed on MAGIC imputed sets of variable genes, as described previously (Brulois, 2020; Xiang *et al*., 2020). Pericytes were removed from this analysis, as too few of these cells were present in the adult dataset to generate meaningful conclusions. Three-dimensional visualization of tSPACE principal components (tPC) 1-3 demonstrated that embryonic cells broadly clustered together and away from adult MSC subsets (Fig. 5C). Nevertheless, two clear connections were observed between embryonic and adult colonic MSC (Fig. 5C, arrow heads). The first was a direct and distinct connection between embryonic clusters 0 and 2 and adult SMC, while the second was a connection between embryonic clusters 4 (mesothelial cells) and 5 to adult CD81^+^ FB and to a lesser extent adult CD90^+^ FB (Fig. 5C). Supporting the idea that the mesothelium gives rise to SMC and some FB in the intestinal serosa and muscle layers of the intestine (Wilm *et al*., 2005; Rinkevich *et al*., 2012), we found that both mesothelial cells and cluster 5 expressed several genes previously associated with FB progenitors (Bae *et al*., 2011; Dulauroy *et al*., 2012; Castagnaro *et al*., 2013; Driskell *et al*., 2013; Worthley *et al*., 2015; Vallecillo-García *et al*., 2017) (Fig. S5B). We thus selected mesothelial cells as a tSPACE trajectory starting point for pseudotime analysis (Fig. 5D). This demonstrated a pseudotime trajectory of mesothelial cells to adult SMC via embryonic clusters 0 and 2 and from mesothelial cells and cluster 5 to adult CD81^+^ FB (Fig. 5D, arrows).

**Figure 5.**
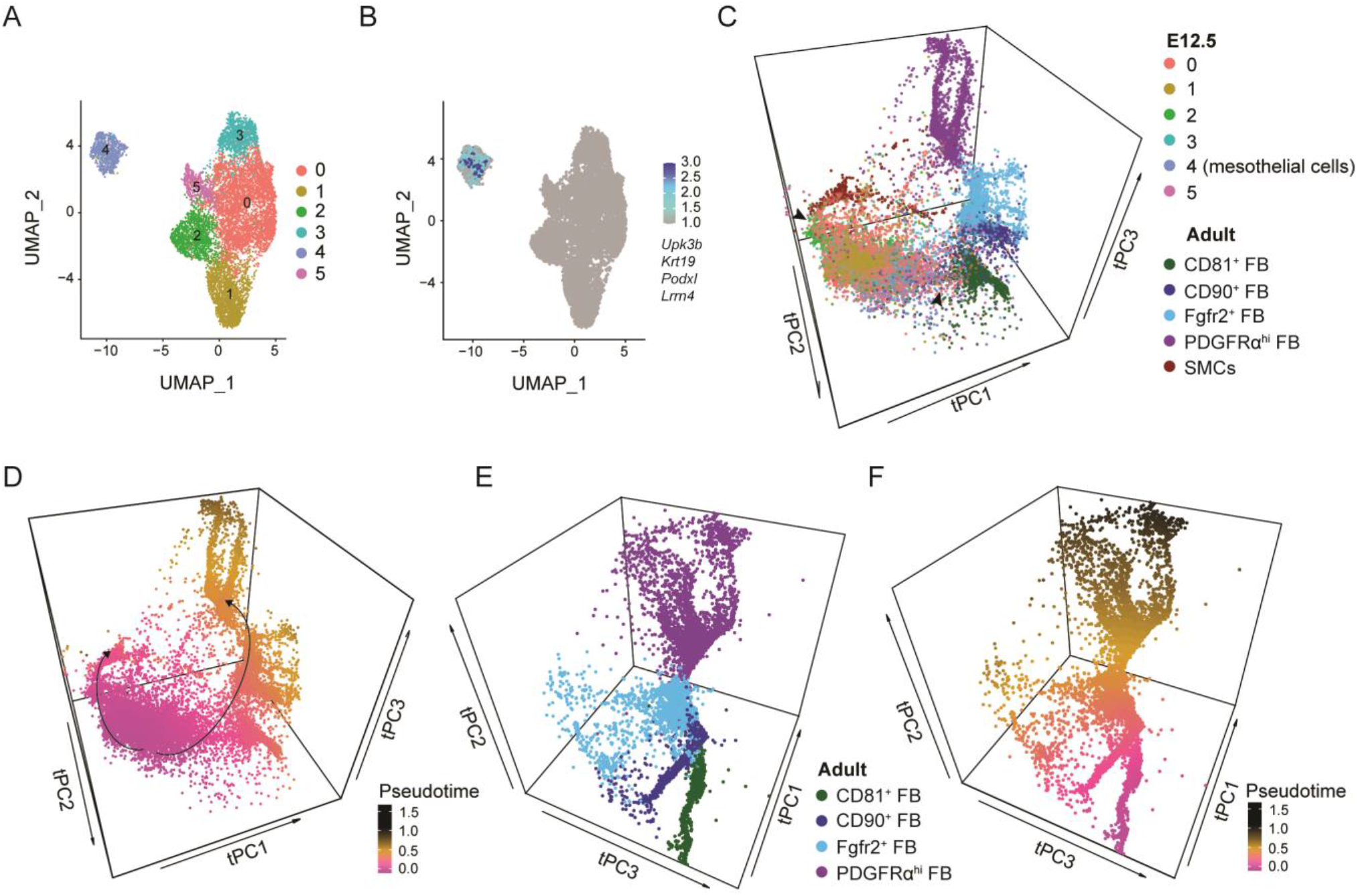
Trajectory analysis indicates that adult intestinal MSC subsets derive from embryonic *Gli1^+^* mesothelial cells. (**A**) UMAP dimensionality reduction of scRNA-seq data colored by Louvain clustering from FACS purified Itgβ1^+^ MSC from the colon of embryonic day E12.5 mice. Data are from 9632 single cells from 2 pooled experiments using 3-5 embryonic colons/experiment, with an average of 2521 genes/cell. (**B**) UMAP of E12.5 large intestinal Itgβ1^+^ MSC overlaid with expression of the indicated mesothelium associated genes. (**C** and **D**) tSPACE principal component analysis (tPC 1-3) projection of pooled adult colonic and E12.5 large intestinal MSC. (**C**) Clusters are color coded as in (**A**) for embryonic clusters or as in Fig. 1B for adult clusters. Arrow heads indicate connections between embryonic and adult clusters. (**D**) Pseudotime analysis using averaged values of the 9 trajectories with starting point in mesothelial cells superimposed on tPC 1-3. (**E** and **F**) tSPACE projections of adult colonic MSC in tPC1-3. (**F**) Pseudotime analysis superimposed on (**E**) using averaged values of the 215 trajectories starting in CD81^+^ FB. See also Figure S5.

Interestingly, rather than branching immediately into distinct FB subsets, adult CD81^+^ FB connected directly to CD90^+^ FB that then connected to Fgfr2^+^ FB and finally to PDGFRα^hi^ FB (Fig 5C), and tSPACE analysis of adult FB showed a similar linear connection between FB subsets (Fig. 5E). As adventitial CD81^+^ FB have been suggested to contain FB precursors in adults (Buechler *et al*., 2021), we used them as the starting population for a new pseudotime analysis, which again indicated a linear trajectory from adult CD81^+^ FB via CD90^+^ FB and Fgfr2^+^ FB to PDGFRα^hi^ FB (Fig. 5F). This conclusion was further supported when we overlaid our trajectory on to the DEG genes for clusters generated by a recent pseudotime analysis of mouse tissue FB, which has suggested a developmental trajectory from *Pi16*^+^ precursors through a population of *Col15α1*^+^ FB that eventually gives rise to mature tissue specific FB that include *Fbln1*^+^and then *Bmp4*^+^ FB in the intestine (Buechler *et al*., 2021). This analysis showed that the *Pi16*^+^, *Col15α1*^+^, *Fbln1*^+^ and *Bmp*4^+^ FB clusters defined by Buechler *et al* broadly overlapped with our colonic CD81*^+^,* CD90*^+^,* Fgfr2*^+^* and PDGFRα^hi^ FB subsets, respectively (Fig. S5C). Collectively, these results suggest that adult MSC subsets originate from the embryonic *Gli1^+^* mesothelium, with adult SMC deriving from an embryonic intermediate distinct from that which gives rise to adult FB. In addition, these results suggest that adult FB subsets arise sequentially from CD81^+^ FB.

Each FB cluster had an extended appearance in tSPACE, with groups of cells streaming outwards from a central core (Fig 5E). To determine what processes might underlie this appearance, we assessed differences in gene expression between the start (core) and end (tip) of each FB cluster (Fig. S5D). While cells at the tip and core of the clusters had similar read counts and detected genes (Fig. S5E), those at the core of each subset expressed 108-121 genes at significantly higher levels than those at the tip, while tip cells expressed no or few (0-2) genes at a significantly higher level. Enrichr based analysis (GO biological process 2018 (Ashburner *et al*., 2000)) of the genes expressed preferentially by cells at the core demonstrated that 9 of the top 10 pathways were shared across FB subsets (Fig. S5F). These included processes involved in positive regulation of transcription, responses to cytokines, and responses to unfolded proteins, with the overwhelming majority of these genes being shared by the core cells in all the FB subsets (Fig. S5G). Together these results suggest that the tip cells within each FB subset are more quiescent than their core counterparts and hence may be more highly differentiated.

### Colonic PDGFRα^hi^ FB consist of three transcriptionally distinct clusters originating from Fgfr2^+^ FB

tSPACE analysis of adult FB subsets indicated that PDGFRα^hi^ FB originated from Fgfr2^+^ FB and then separated into three branches (Fig. 5E and F). To validate the idea that Fgfr2^+^ FB act as precursors of PDGFRα^hi^ FB we performed RNA velocity analysis (La Manno *et al*., 2018) focusing on these subsets, which confirmed the directionality from Fgfr2^+^ FB to PDGFRα^hi^ FB (Fig. 6A). Re-clustering of only PDGFRα^hi^ FB uncovered three clusters that diverged along the three trajectory branches (Fig. 6B) that could be distinguished based on expression of *Cd9* and *Cd141* (thrombomodulin (Thbd)) (Fig. 6C). This generated clusters of CD9^hi^CD141^-^, CD9^lo^CD141^+^ and CD141^int^ cells, all of which expressed the “telocyte” marker, *Fox1l* (Fig. S6A) (Shoshkes-Carmel *et al*., 2018). Consistent with these findings, flow cytometric analysis of colonic PDGFRα^hi^ FB identified distinct clusters of CD9^hi^CD141^-^ and CD9^lo^CD141^+^ cells, together with CD9^-^ cells that expressed heterogeneous levels of CD141 and which we referred to as CD141^int^ FB (Fig. 6D). Analysis of the top DEG between these populations demonstrated that CD9^hi^CD141^-^ cells expressed the highest levels of *Nrg1*, *Fgf7, Il1rl1* (ST2 (IL33 receptor)) and *Ptgs2*, that CD9^lo^CD141^+^ cells expressed high levels of fibrosis-associated *Aspn* (Asporin), *Il11ra1* and *Cxcl12*, while CD141^int^ cells expressed high levels of *Cxcl10*, *Ly6c1*, *Adamdec1*, *Wnt4a* and *Plpp3* (Fig. S6B). The CD9^lo^CD141^+^ cells, and to a lesser extent the CD141^int^ cells, expressed mRNA and protein for αSMA (Fig. S6C), a marker of myofibroblasts but not “telocytes”. Immunohistochemical staining for PDGFRα and αSMA showed that αSMA^+^PDGFRα^hi^ cells localize preferentially to the isthmus area just above colonic crypts, while αSMA^-^PDGFRα^hi^ cells aligned directly underneath the epithelium at the top and bottom of crypts (Fig. 6F). The CD9^lo^CD141^+^, CD9^hi^CD141^-^ and CD141^int^ FB also differentially expressed several epithelial support genes (Fig. 6G), suggesting that these populations may play distinct roles in supporting the epithelium at different stages of its development. Thus, adult colonic subepithelial PDGFRα^hi^ FB consist of spatially and transcriptionally distinct clusters that derive from Fgfr2^+^ FB.

**Figure 6.**
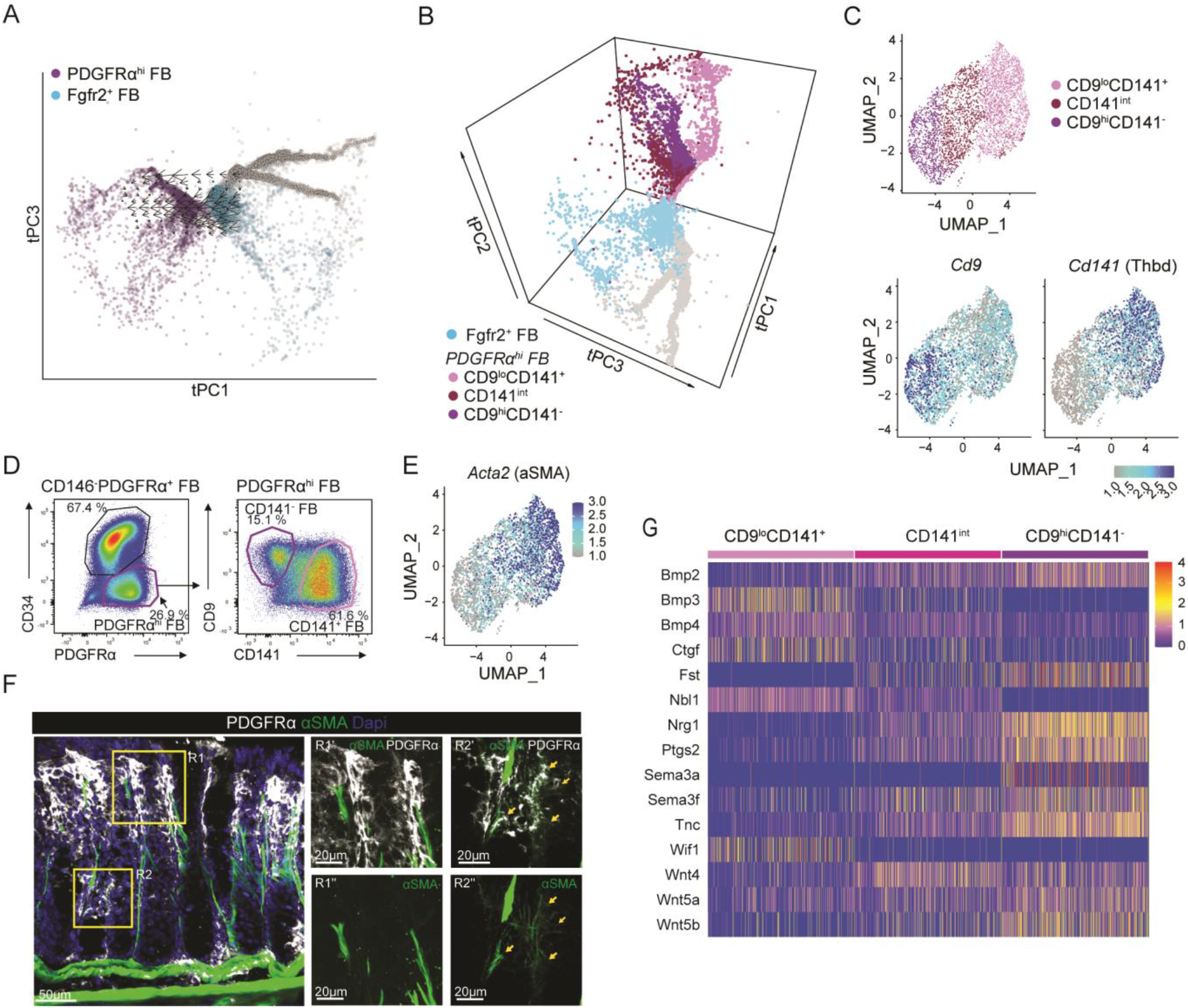
Subepithelial PDGFRα^hi^ FB consist of three transcriptionally distinct clusters originating from Fgfr2^+^ FB. (**A**) tSPACE projection of adult colonic MSC in tPC1 and 3 highlighting Fgfr2^+^ FB and PDGFRα^hi^ FB overlaid with RNA Velocity. (**B**) tSPACE projection of colonic MSC in tPC1-3, highlighting Fgfr2^+^ FB and three PDGFRα^hi^ FB clusters. (**C**) UMAP dimensionality reduction of re-clustered colonic PDGFRα^hi^ FB (top panel), with *Cd9* (bottom left panel) and *Cd141* (bottom right panel). (**D**) Representative flow cytometric analysis of CD9 and CD141 expression by colonic PDGFRα^hi^ FB. Representative plots from 2 experiments with 3 mice/experiment. (**E**) *Acta2* (αSMA) gene expression projected onto UMAP of colonic PDGFRα^hi^ FB. (**F**) Immunohistochemical staining of colonic tissue for indicated antigens. R1’,” and R2’,” represent magnifications of R1 and R2 quadrants (yellow squares) on left image. Results are representative stains from 3 experiments with 3 mice/experiment. Arrows indicate αSMA^+^PDGFRα^hi^ FB. (**G**) Heatmap showing scaled transcription levels (integrated data) of significantly (p<0.05) differentially expressed epithelial support genes between the PDGFRα^hi^ FB clusters. See also Figure S6.

## Discussion

Recent studies have demonstrated considerable heterogeneity within the intestinal LP MSC compartment (Degirmenci *et al*., 2018; Kinchen *et al*., 2018; Bahar Halpern *et al*., 2020; Hong *et al*., 2020; McCarthy *et al*., 2020; Roulis *et al*., 2020) and suggested non-redundant roles for MSC subsets in intestinal homeostasis (Bahar Halpern *et al*., 2020; David *et al*., 2020; Hong *et al*., 2020; McCarthy *et al*., 2020; Wu *et al*., 2021), inflammation (West *et al*., 2017; Kinchen *et al*., 2018; Smillie *et al*., 2019) and cancer (Roulis *et al*., 2020). As these studies have largely focused on single regions of the intestine (Degirmenci *et al*., 2018; Kinchen *et al*., 2018; Smillie *et al*., 2019; Bahar Halpern *et al*., 2020; David *et al*., 2020; Hong *et al*., 2020; McCarthy *et al*., 2020; Roulis *et al*., 2020; Wu *et al*., 2021), it has been unclear whether there are regionally circumscribed differences in the composition of LP MSC subsets along the length of the intestine. By performing scRNA-seq analysis of small intestinal and colonic LP MSC from the same mice, we show here that both locations contain similar LP MSC subsets and that their pattern of expression of epithelial support genes by LP MSC subsets is largely conserved between these sites. Bulk RNA-seq analysis of sorted PDGFRα^hi^ subepithelial FB and interstitial Fgfr2^+^ FB confirmed that these subsets expressed distinct arrays of epithelial support genes irrespective of the tissue. However, both PDGFRα^hi^ and Fgfr2^+^ FB expressed higher levels of many epithelial support genes in the colon compared with their small intestinal counterparts, indicating a greater role for these FB in sustaining epithelial integrity in the colon. Consistent with this idea, WNT secretion by *Gli1*-expressing MSC is essential for homeostasis of the colonic epithelium (Degirmenci *et al*., 2018; Karpus *et al*., 2019; David *et al*., 2020), but this is not the case in the small intestine where Paneth cells represent a major source of WNTs (Sato *et al*., 2011).

Among the pathways significantly upregulated in colonic FB subsets compared with those in small intestine were *TGFβ regulation of extracellular matrix expression*, *epidermal growth factor receptor (EGFR1) signaling*, *brain derived neurotrophic factor (BDNF) signaling pathway* and *thyroid stimulating hormone (TSH) regulation of gene expression*. The relevance of these pathways in colonic versus small intestinal homeostasis remains to be determined. In contrast, few pathways were selectively upregulated in small intestinal FB subsets. Among these, several genes encoding chemokines, cytokines and cytokine receptors were significantly overexpressed by the Fgfr2^+^ FB population, suggesting that this subset may be involved in immune functions in the small intestine, but not in the colon. Interestingly, small intestinal FB also expressed higher levels of enzymes involved in vitamin A metabolism, consistent with previous findings that some small intestinal FB display aldehyde dehydrogenase activity and that there is increased retinoic acid receptor signaling in the small compared with the large intestine (Jaensson-Gyllenbäck *et al*., 2011; Vicente-Suarez *et al*., 2015). Collectively, these findings indicate that the local microenvironment plays a crucial role in regulating the transcriptional profile and specialization of intestinal FB in different regions of the intestine. The nature of the relevant factors and their importance in local homeostasis awaits further study.

Consistent with the idea that they may provide niche-specific support for local cells, the subsets of intestinal FB were located within distinct regions of the gut wall. As others have shown (Eyden, Curry and Wang, 2011; Kurahashi *et al*., 2013; Shoshkes-Carmel *et al*., 2018), we found that PDGFRα^hi^ FB directly underlie the intestinal epithelium in both the small and large intestine. In contrast to an earlier report that small intestinal CD81^+^ FB lie solely within the submucosa (Thomson *et al*., 2018), we found these cells within both the submucosa and surrounding larger vessels deep in the mucosa, consistent with more recent studies (Hong *et al*., 2020; McCarthy *et al*., 2020). In addition, we demonstrate that CD81^+^ FB are found in similar locations in the small intestine and colon. CD81^+^ FB play an essential role in maintaining the epithelial stem cell niche in the small intestine, partly through their selective expression of the BMP antagonist, gremlin-1 (McCarthy *et al*., 2020). Consistent with this, our scRNA-seq analysis demonstrated that CD81^+^ FB were the major source of *Grem1* in the small intestine, although CD81^+^ FB, CD90^+^ FB and Fgfr2^+^ FB all expressed *Grem1* in the colon, indicating potential redundancy between these subsets in supporting the colonic epithelial stem cell niche.

Less is known regarding the location of the intestinal PDGFRα^lo^ FB subsets that do not express CD81. Our immunohistochemical analysis demonstrated that Fgfr2^+^ FB were located preferentially within the villus core towards the villus tip in the small intestine, while those in the colon were located between crypts. These findings are consistent with work on *Fgfr2-* mCherry reporter mice that suggested the Fgfr2^+^ cells represent interstitial FB (Roulis *et al*., 2020). Using PPARγ expression as a surrogate marker, we could show that CD90^+^ (PPARγ^+^) FB were located near the base of colonic crypts, but were unable to do this in the small intestine, as we failed to identify a specific marker for this population. However, small intestinal Igfbp5^+^ FB and colonic CD90^+^ FB resembled one another transcriptionally and Igfbp5^+^ FB were the only CD34^+^CD81^-^ FB subset in small intestine. As the cells with this phenotype were located primarily around small intestinal crypts, these results indicate that CD90^+^ FB and Igfbp5^+^ FB appear to represent a peri-cryptal population of FB in the colon and small intestine, respectively. Collectively, our findings confirm and extend previous work on the localization of intestinal FB subsets, highlighting their distinct transcriptional profiles and complex spatial organization within the mucosa.

The mesothelium is an epithelial monolayer that lines body cavities and internal organs, including the serosal surface of the intestine (Winters and Bader, 2013). Mesothelial cells undergo epithelial-mesenchymal transition (EMT) and can give rise to both SMC and FB in response to injury in a number of tissues including the intestine (Miyoshi *et al*., 2012; Rinkevich *et al*., 2012; Koopmans and Rinkevich, 2018). During ontogeny, the intestinal mesothelium is a source of precursors for SMC in the intestinal muscle layers and vasculature (Wilm *et al*., 2005), as well as for uncharacterized FB in the outer serosa of the intestine (Rinkevich *et al*., 2012). In contrast, the origin(s) of the MSC subsets in the adult small intestine and colon LP has remained unclear. Here we used intestinal transplantation and lineage-tracing approaches to demonstrate that all adult small intestine and colon LP MSC subsets derive from *Gli1*-expressing progenitors present in the E12.5 intestine. At that time point, *Gli1* expression was restricted to mesothelial cells and an embryonic population of CD34^-^PDGFRα^+^ FB. scRNA-seq analysis showed that both mesothelial cells and a minor cluster of cells within the CD34^-^PDGFRα^+^ FB population expressed markers previously associated with FB progenitors, while tSPACE analysis suggested a direct trajectory connection between these two populations. These results provide strong evidence that the mesothelium is a source of FB precursors during early intestinal development and that these are capable of giving rise to all adult LP MSC subsets. Interestingly, while also orignating from embryonic mesothelium, smooth muscle cells (SMC) in adult LP developed along an embryonic trajectory that was distinct from that of adult FB in tSPACE. Thus, SMC and FB in the steady state adult LP appear to represent independent lineages that are specified during development.

tSPACE analysis revealed direct connections between embryonic MSC clusters and adult CD81^+^ FB, suggesting that adult FB subsets arise from CD81^+^ FB, rather than from distinct populations of intermediates that develop in the embryo. Consistent with this idea, CD81^+^ FB locate in the submucosa and in the adventitia surrounding larger vessels at the base of the mucosa, an anatomical niche that contains MSC progenitors in other tissues (Sidney *et al*., 2014; Díaz-Flores *et al*., 2015; Kramann *et al*., 2015, 2016; Sitnik *et al*., 2016; Benias *et al*., 2018; Merrick *et al*., 2019). Furthermore, lineage tracing of *Grem1^+^* FB in the adult small intestine has identified progenitors that give rise to subepithelial FB along the entire crypt-villus axis, in a process that is relatively rapid during the perinatal period, but takes around a year to be completed in adult intestine (Worthley *et al*., 2015). In support of this idea, we found that *Grem1* expression is largely restricted to CD81^+^ FB in the small intestine.

While early trajectory analysis suggested a bifurcation downstream from CD81^+^ FB (Kinchen *et al*., 2018), our tSPACE, pseudotime and Velocity analyses suggested that adult colonic CD81^+^ FB connected in a linear direction to adult CD90^+^ FB, then to Fgfr2^+^ FB and finally to PDGFRα^hi^ FB. Although we used different markers, our findings are consistent with those recently published by Buechler *et al* who suggested that adventitial *Pi16*^+^ FB give rise first to *Col15α1*^+^ FB and then to tissue specific FB clusters that in the intestine included *Fbln1*^+^ FB and subsequently *Bmp4*^+^ FB (Buechler *et al*., 2021). Overlaying these clusters on our tSPACE trajectories demonstrated that the *Pi16*^+^, *Col15α1^+^* and *Fbln1*^+^ FB most closely resembled the CD81^+^, CD90^+^ and Fgfr2^+^ FB we found in colon, while the *Bmp4*^+^ FB were similar to our colonic PDGFRα^hi^ FB. The trajectory from CD81^+^ FB also correlates with the basal to apical localization of the downstream subsets in the colonic LP, indicating that this process may reflect factors present in distinct microenvironmental niches. Despite this strong evidence for linear differentiation from a single precursor, it should be noted that FB show evidence of slow turnover in the adult intestine (Worthley *et al*., 2015; Kinchen *et al*., 2018; Bahar Halpern *et al*., 2020) and we cannot exclude the possibility that each FB subset might self-renew *in situ*.

Recent scRNA-seq studies have suggested that colonic subepithelial PDGFRα^hi^ FB are heterogeneous (Kinchen *et al*., 2018; Roulis *et al*., 2020) and here we found that PDGFRα^hi^ FB diverged into 3 clusters, which we could define as CD9^lo^CD141^+^, CD9^hi^CD141^-^ and CD141^int^ FB. The CD9^lo^CD141^+^ FB are likely related to the PDGFRα^hi^ FB sub-cluster S2a defined by Kinchen *et al*, as they expressed high levels of *Cxcl12*, while CD9^hi^CD141^-^ FB expressed high levels of *Nrg1* and so are likely related to the PDGFRα^hi^ FB subcluster S2b (Kinchen *et al*., 2018). Although all three clusters expressed the telocyte marker *Foxl1* (Shoshkes-Carmel *et al*., 2018), CD9^lo^CD141^+^ FB and to a lesser extent CD141^int^ FB, expressed *Acta2*, coding for α−smooth muscle actin (αSMA), a marker often associated with myofibroblasts. αSMA^+^PDGFRα^hi^ FB were located directly underneath the epithelium approximately half way up colonic crypts, suggesting that CD9^lo^CD141^+^ and CD9^hi^CD141^-^ subepithelial FB localise within distinct regions along the crypt axis. Of the three PDGFRα^hi^ FB clusters, CD9^lo^CD141^+^ FB expressed the highest levels of the WNT antagonists *Wif1*, *Bmp3* and *Bmp4*. Therefore we speculate that the location of CD9^lo^CD141^+^ FB half way up colonic crypts allows them to promote the terminal differentiation of epithelial cells as they migrate up the crypt (Qi *et al*., 2017). In contrast, CD9^hi^CD141^-^ FB expressed high levels of top of crypt-associated non-canonical *Wnt4, Wnt5a* and *Wnt5b* (Gregorieff *et al*., 2005; Kosinski *et al*., 2007) and *Tenascin C (Tnc)* (Probstmeier, Martini and Schachner, 1990; Bernier-Latmani *et al*., 2015) and base of crypt-associated *Ptgs2* (the gene encoding COX-2) (Stzepourginski *et al*., 2017; Roulis *et al*., 2020), *Sema3a* (Karpus *et al*., 2019). Thus, our findings indicate that each of the three PDGFRα^hi^ FB subsets may play distinct roles in colonic epithelial homeostasis.

In conclusion, our study provides a comprehensive mapping of intestinal MSC diversity, location and epithelial support function and highlights a central role for location along the intestinal length in regulating transcriptional profile and functional specialization. We also show that all adult MSC derive from *Gli1*-expressing embryonic mesothelial cells and we propose there is a linear developmental relationship between adult FB subsets that culminates in the development of a heterogeneous group of subepithelial PDGFRα^hi^ FB. Together our findings provide key insights into MSC diversity, development, function and interrelationships with relevance to intestinal development and homeostasis.

## Acknowledgements

We thank Dr. J. Vandamme (DTU, Denmark) for performing scRNA-seq and library preparation, Dr. A.L Joyner (Memorial Sloan-Kettering Cancer Center) for providing *Gli1*-EGFP mice, Dr. S. Milling (University of Glasgow University, UK) for providing laboratory space and materials for experiments involving *Ackr4*^tmlCcbl1^ mice and Dr. R. Gentek (Edinburgh University, UK) for advice regarding embryonic lineage tracing. The SNP&SEQ Platform is part of the National Genomics Infrastructure (NGI) Sweden and Science for Life Laboratory. The SNP&SEQ Platform is also supported by the Swedish Research Council and the Knut and Alice Wallenberg Foundation. This work was supported by grants awarded to W.W.A. from the Lundbeck foundation (R155-2014-4184), Denmark, and the Gut Cell Atlas, an initiative funded by the Leona M. and Harry B. Helmsley Charitable Trust, US.

## Author contributions

The study was designed by S.I.P, S.S., and W.W.A. Experiments in Denmark were performed by S.I.P., S.S. U.M and J.J., the grafting experiments were performed by K.W. and S.I.P, experiments in Glasgow were performed by S.I.P. and A.T.A. with support from R.J.B.N., bioinformatics analyses was performed by L.W. with support from K.N., K.F.B, E.C.B and S.B. A.M provided valuable intellectual input throughout. The manuscript was written by S.I.P. and W.W.A after input from all authors.

## METHODS

### RESOURCE AVAILABILITY

#### Lead Contact

Further information and requests for resources and reagents should be directed to and will be fulfilled by the Lead Contact, Dr WW Agace.

#### Materials Availability

This study did not generate new unique reagents or mouse strains.

#### Data and Code Availability

Single-cell RNA-seq and bulk RNA-seq data has been deposited at NCBI GEO and are publicly available as of the date of publication. Accession numbers will be listed when published. Microscopy data reported in this paper will be shared by the lead contact upon request. This paper does not report original code. Any additional information required to re-analyze the data reported in this paper is available from the lead contact upon request.

### EXPERIMENTAL MODEL AND SUBJECT DETAILS

#### Mice

*Gli1*^tm3(cre/ERT2)Alj^ (*Gli1*-CreER^T2^, 007913 Jackson laboratories), B6.129X1-Gt(ROSA)26Sor^tm1(EYFP)Cos^/J (*R26R*.EYFP, 006148 Jackson laboratories), *Gli1*-EGFP (Garcia *et al*., 2010) and EYFP mice (obtained by crossing *R26R*.EYFP with the relevant Cre mice) were bred and maintained at the Bio-Facility animal house (Technical University of Denmark). C57BL/6Nrj mice were purchased from Janvier Labs (Le Genest-Saint-Isle, France). *Ackr4*^tmlCcbl1^ mice (*Ackr4*.EGFP) (Heinzel, Benz and Bleul, 2007) were bred and maintained in the Central Research Facility, Glasgow University. Adult mice were used between 5.5 and 12w of age. Mice of both genders were used in all experiments and littermates were used as controls. All experiments were approved by the Danish Animal Experiments Inspectorate, or with ethical approval under a Project Licence from the the UK Home Office.

## METHOD DETAILS

### Kidney grafting

EYFP male mice were mated overnight with C57BL/6Nrj females and the following morning was defined as gestational day 0.5 (E0.5). Pregnant dams were sacrificed at E12.5 and small and large intestine were dissected from embryos under a stereo microscope (VWR). Adult WT mice were anaesthetized by i.p injection of Ketaminol Vet. (100mg/kg, MSD animal health) and Rompun Vet. (10mg/kg, Bayer) and were injected subcutaneously with Bupaq (0.1mg/kg, Richter Pharma). Washed embryonic intestine was transplanted under the kidney capsule of anesthetized recipients as described previously (Ferguson, Parrott and Connor, 1972). Recipients were sacrificed at the time points indicated and grafts were dissected and cut into pieces prior to cell isolation as described below.

### *In vivo* lineage tracing

*Gli1*-CreER^T2^ male mice were mated overnight with *R26R*.EYFP females and the following morning was defined as gestational day E0.5. At E11.5, pregnant dams were injected i.p. with 4-hydroxytamoxifen ((4-OHT), 1.6 mg, Sigma) and progesterone (0.8 mg, Sigma) in 160μl PBS with 25% Kolliphor (Sigma)/25% ethanol (Fischer Scientific). Small and large intestine were isolated from embryos or weaned offspring at the time points indicated.

### Cell isolation

Intestinal cell suspensions were generated as described previously (Schulz *et al*., 2009) with minor changes. Briefly, washed intestinal tissue was opened longitudinally and Peyer’s patches removed. For scRNA-seq and bulk RNA-seq experiments on adult intestine, muscularis externa was stripped away using tweezers. Tissues were cut into 0.5-1 cm pieces and epithelial cells removed by 3 consecutive rounds of incubation in HBSS supplemented with HEPES (15mM), sodium pyruvate (1mM), penicillin/streptomycin (100 U/mL), gentamycin (0.05 mg/mL), EDTA (2mM) (all Invitrogen) and FCS (2.5%) (Sigma), for 15 min at 37°C with constant shaking at 350 rpm. After each incubation, samples were shaken for 10 sec and medium containing epithelial cells and debris was discarded. For colonic tissues, DL-dithiothreitol (5mM) (Sigma) was added at the first incubation step. Remaining tissue pieces were digested with collagenase P (0.6U/mL, Sigma) or with Liberase TM (0.325U/mL, Roche) and DNAse I (31 μg/mL, Roche) in R10 medium (RPMI 1640, sodium pyruvate (1mM), HEPES (10 mM), penicillin/streptomycin (100 U/mL), gentamycin (0.05mg/mL), and 10% FCS) for up to 30 min at 37°C with constant shaking at 550 rpm (small intestine) or with a magnetic stirrer and at 280 rpm (large intestine). For bulk RNA-seq cells were treated with ACK lysing buffer (Gibco) to lyse red blood cells prior to sorting. For isolation of cells from embryonic intestine, tissues were digested directly for 30 min at 37°C in Eppendorf tubes with constant shaking at 900 rpm. The resulting cell suspensions were filtered through a 70 μm filter and washed in MACS buffer (phosphate buffered saline (PBS) with 3% FCS and 2 mM EDTA) twice prior to subsequent analyses.

### Flow cytometry and cell sorting

Cell suspensions were stained with fluorochrome labelled primary antibodies (see table 1) in Brilliant stain buffer (BD Biosciences) for 30 min on ice. Flow cytometry was performed on an LSR Fortessa II (BD Biosciences), FACSAria Fusion (BD Biosciences), or FACSMelody (BD Biosciences) and analysed with FlowJo software (TreeStar). Dead cells were identified by staining with either 7-AAD (eBioscience) or Zombie UV fixable viability dye (BD Biosciences) and cell doublets were excluded on the basis of FSC-A/FSC-H. For intracellular staining, cells were stained for surface antigens, fixed with FoxP3 Staining Buffer set (eBioscience) and stained for αSMA in FoxP3 Permeabilization buffer (eBioscience). After washing, cells were stained with antibodies to surface antigens not compatible with fixation according to the manufacturer’s instructions.

**Table 1:**
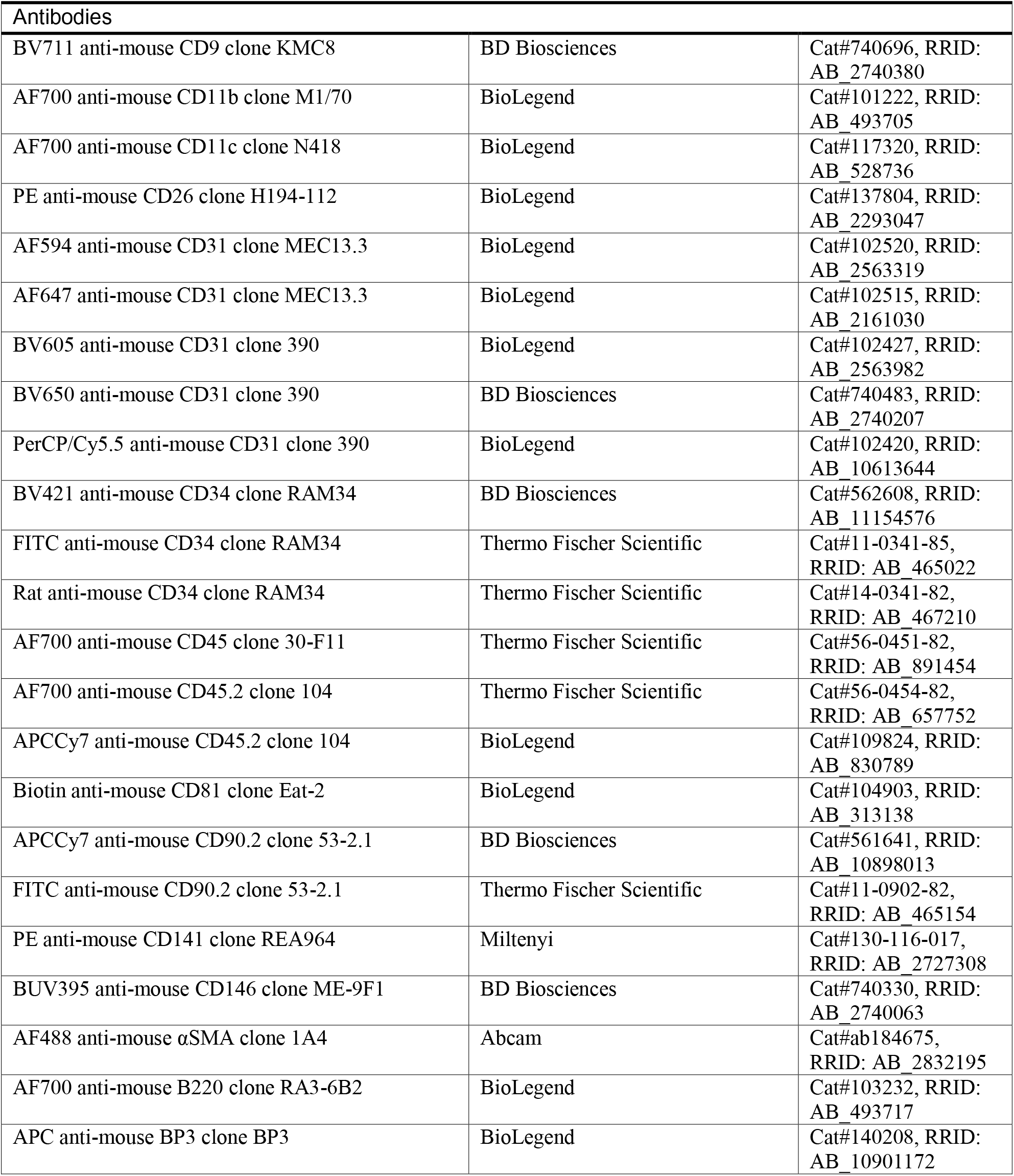
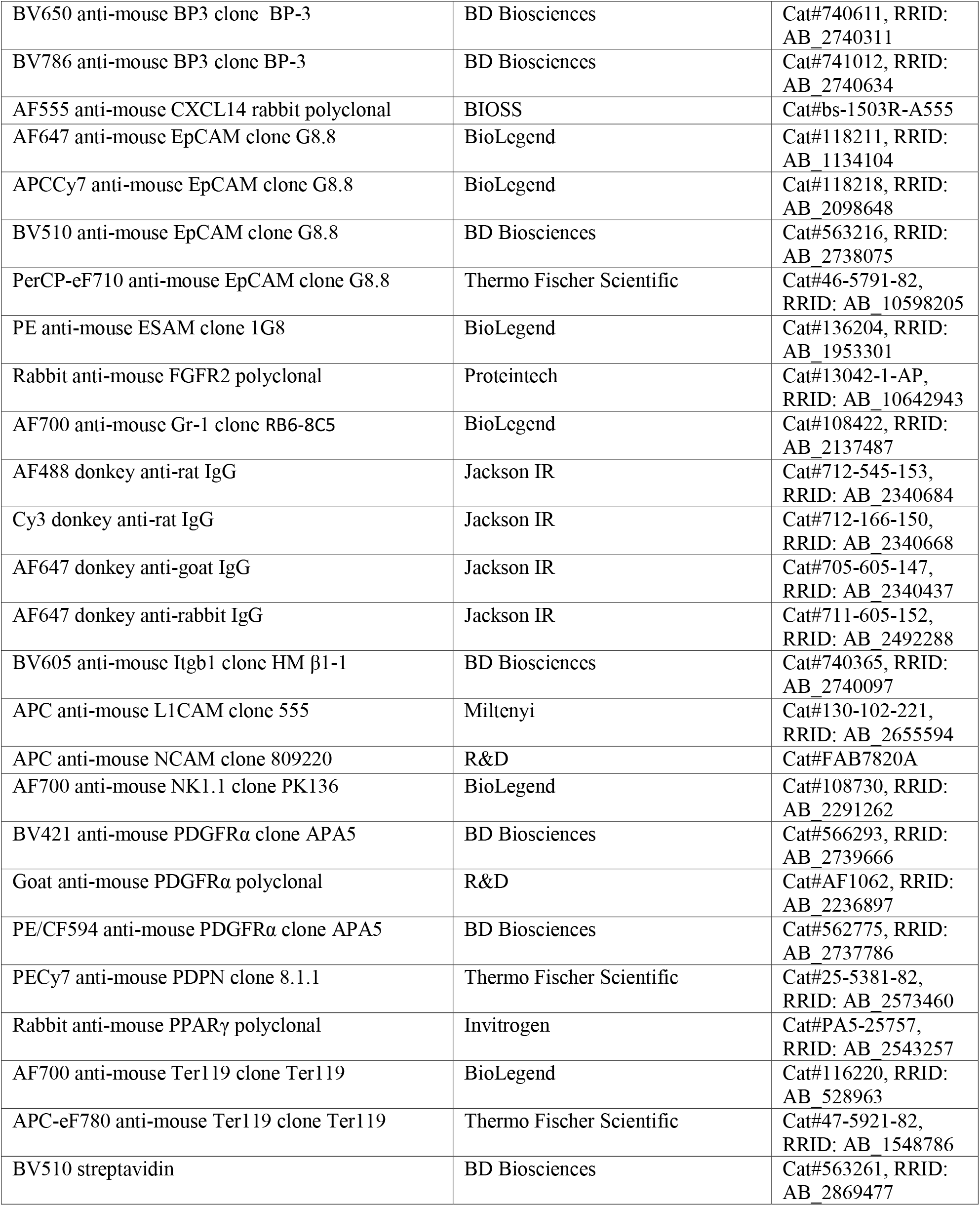
Antibodies

### Immunohistochemistry

Tissues were fixed in paraformaldehyde (4%, PFA) and sectioned (70 μm) using a Vibratome (Leica VT12000S). Sections were incubated in PBS containing BSA (1%) and Triton-X100 (0.3%) for 1 hour at room temperature (RT) to block non-specific staining and incubated with fluorochrome conjugated or unconjugated primary antibodies (see Table S1) overnight at 4⁰C. After washing with PBS containing Triton-X100 (0.3%), tissues were incubated with secondary antibodies at RT for 2-4 hours (see table 1). For detection of CD81, staining with biotinylated anti-CD81 was enhanced using the Biotinyl Tyramide kit (Perkin Elmer) according to the manufacturer’s instructions after blocking of endogenous biotin using Streptavidin/Biotin Blocking kit (Invitrogen). Endogenous peroxidase was inactivated by incubating tissues with 3% H_2_O_2_ for 30 min at RT before incubation with streptavidin-horse radish peroxidase (HRP). Sections were analysed under 40x magnification using a Zeiss LSM 710 confocal microscope and the images were processed using Zeiss Zen and Imaris software. For histological analysis of kidney grafts, tissue pieces were fixed in 4 % paraformaldehyde for 8 h and paraffin-embedded sections were stained with hematoxylin and eosin.

### Library Preparation and sequencing

#### Single cell RNA-seq

Sorted cells were washed in cold PBS containing bovine serum albumin (0.04%), counted and diluted to the desired concentration following 10X Genomics guidelines (10x Genomics, CG000053_CellPrepGuide_RevC). ScRNA-seq libraries were prepared according to the manufacturer’s instructions using Chromium Single Cell 3′ Library & Gel Bead Kit v3 (10x Genomics, PN-1000092) or 5’ kit Chromium Single Cell 5′ Library & Gel Bead Kit (10x Genomics, PN-1000006) and Chromium Chip B Single Cell Kit (PN-1000074) with the Chromium Controller & Next GEM Accessory Kit (10x Genomics, PN-120223). In brief, single cells, reverse transcription reagents, Gel Beads containing barcoded oligonucleotides, and oil were combined on a microfluidic chip to form Gel Beads in Emulsion (GEMs). Individual cells were lysed inside the GEMs and the released poly-A transcripts were barcoded with an Illumina R1 sequence, a 10X barcode and a Unique Molecular Identifier (UMI) during reverse transcription (RT). After RT, GEMs were broken, barcoded cDNA was purified using Dynabeads MyOne silane (10x Genomics, PN-2000048) and amplified by Polymerase Chain Reaction (PCR). Amplified cDNA were cleaned up with SPRIselect Reagent kit (Beckman Coulter, B23318). Indexed sequencing libraries were constructed by enzymatic fragmentation, end-repair and A-tailing, before a second and final PCR amplification using the Chromium i7 Sample Index (10x Genomics, PN-220103), introducing an Illumina R2 sequence, a unique sample index (allowing multiplex sequencing) and P5/P7 Illumina sequencing adaptors to each library. Library quality control and quantification were performed using a KAPA Library Quantification Kit for Illumina Platforms (Kapa Biosystems, KK4873) and the 2100 Bioanalyzer equipped with a High Sensitivity DNA kit (Agilent, 5067-4626). Muliplexed libraries were pooled and sequenced either by NextSeq 500/550 High Output v2.5 kit (150 cycles) at the Center of Excellence for Fluorescent Bioanalytics (KFB, University of Regensburg, Germany) or by Novaseq 6000 S1 or S2 (200 cycles) at the SNP&SEQ Technology Platform (Uppsala, Sweden) with the following sequencing run parameters: Read1 - 28 cyles; i7 index - 8 cycles; Read2 – 126 cycles at a depth of at least 100M reads/sample.

#### Bulk RNA-seq

Cells were sorted into RLT buffer and total RNA was isolated using the RNeasy Micro kit. Extraction was performed according to the manufacturer’s protocol, with an on-column DNAse digestion step added after the first wash buffer step. RNA quality and quantity were measured using the 2100 BioAnalyzer equipped with RNA6000 Pico chip (Agilent Technologies). RNA was subjected to whole transcriptome amplification using Ovation RNA-Seq System V2 kit and amplified cDNA samples were purified using the MinElute Reaction Cleanup kit. The quantity and quality of the cDNA samples were measured using the 2100 BioAnalyzer equipped with DNA1000 chip (Agilent technologies) and the Nanodrop (ThermoFisher Scientific). Libraries were constructed with the Ovation Ultralow system V2 kit, following the manufacturer’s instructions. Amplified cDNA (100 ng) was fragmented by sonication using a Bioruptor Pico (Diagenode), sheared cDNA end-repaired to generate blunt ends and ligated to Illumina adaptors with indexing tags followed by AMPure XP bead purification. Library size distribution was evaluated using a 2100 Bioanalyzer equipped with DNA1000 chip (Agilent technologies) and quantified using KAPA library Quantification Kit Illumina platforms (Kapa Biosystems). Libraries were diluted and pooled at equimolar concentration (10 nM final) and sequenced on the Hiseq2500 platform (Illumina) using 50bp single reads (Center for Genomic Regulation, Spain) with a depth of 15-20M reads per sample.

### Computational analysis

#### Single cell RNA-seq

Alignment of scRNA-seq data to reference genome mm10 was performed with CellRanger (version 3.0.2 & 3.1.0) (Dobin *et al*., 2013; Zheng *et al*., 2017). The data was imported into R (version 4.0.1) (R Core Team, 2020) and processed to remove debris and doublets in individual samples by looking at gene, read counts and mitochondrial gene expression. Variable genes were calculated with Seurats FindVariableFeatures function and selection method set to “vst” (version 3.1.5) (Stuart and Satija, 2019). The respective samples and all their overlapping genes were then integrated with anchor integration for Seurat. Cell cycle effects were regressed out with linear regression using a combination of the build-in function in Seurat and scoring gene sets from ccremover per cell (Li, Jun; Barron, 2017) during scaling of the gene expression. The datasets were dimensionality reduced first with PCA and then UMAP and clustered with Louvain clustering all using Seurat. After initial clustering, contaminating cells were removed and an additional round of clustering and dimensionality reduction with UMAP was run on the cells of interest. DEGs were identified using Seurat FindAllMarkers function with the default test setting (non-parameteric Wilcoxon Rank Sum test). Expression of gene modules in the form of published signature gene sets were calculated with AddModuleScore (Seurat) taking the top DEG from telocytes (10 genes), Lo-1 FB (10 genes) and Lo-2 FB (7 genes) reported by McCarthy *et al* (McCarthy *et al*., 2020), the top 10 DEG from FB1-5, MC and SMCs reported by Hong *et al* (Hong *et al*., 2020), and the top 20 DEG from *Pi16^+^*, *Col15a1^+^*, *Fbln1^+^* and *Bmp4^+^* FB reported by Buechler *et al* (Buechler *et al*., 2021). Pearson correlations between datasets were calculated based on average expressions per cluster of overlapping variable genes and plotted with heatmap.2 (version 3.0.3) (Gregory R. Warnes, Ben Bolker, Lodewijk Bonebakker, Robert Gentleman, Wolfgang Huber, Andy Liaw, Thomas Lumley, Martin Maechler, Arni Magnusson, Steffen Moeller and Venables, 2020). Heatmaps were constructed with a modified version of Seurats DoHeatmap to allow for multiple grouping variables. Data plotted in expression heatmaps was scaled based on the anchor integrated data. Data imputation was performed per dataset across samples on raw count data with magicBatch (Brulois, 2020) where the affinity matrix used was Seurat’s batch-corrected PCA coordinates. Trajectories and trajectory spaces were determined with tSPACE (Dermadi *et al*., 2020) on the top 2000 imputed variable genes for adult trajectories and top 1000 imputed variable genes for the integrated E12.5 and adult trajectory. GO analysis was performed using GO Biological Process 2018 (Ashburner *et al*., 2000) from Enrichr computational biosystems (Chen *et al*., 2013; Kuleshov *et al*., 2016; Xie *et al*., 2021).

#### Bulk RNA-seq

Raw RNA sequencing data from the 30 samples were pre-processed with TrimGalore (version 0.4.0) and FastQC (version 0.11.2). Pseudo-alignment of reads was performed with Kallisto (version 0.42.5) to obtain RNA expression information. To assess correlations between bulk-seq samples and SC clusters Pearson correlations based on SC variable genes were calculated between the bulk-seq samples and the pseudo-bulk of the SC clusters for the individual tissues and visualized with heatmap.2 (part of gplot package). For all DESeq2 (1.26.0 (Love, Huber and Anders, 2014)) analysis, transcripts identified in less than 3 replicates and at levels below 6 reads were filtered out prior to further analysis. Heatmaps of bulk-seq data expression was created in R with the ComplexHeatmap package (version 2.7.11) and volcano plots with ggplot2 (version 3.3.1). For the comparison between tissues, DEGs were only classified as significant if they had a |log2FC| > 1.5 and adjusted p-value < 0.05. GO analysis was performed using BioPlanet 2019 (Huang *et al*., 2019) from Enrichr computational biosystems (Chen *et al*., 2013; Kuleshov *et al*., 2016; Xie *et al*., 2021).

### Statistical analysis

Statistical significance was determined with a 2-way ANOVA with Benjamini, Krieger and Yekutieli multiple comparisons and performed in Prism software (GraphPad). *p < 0.05, **p < 0.01, ***p < 0.001.

## Supplementary Information

The manuscript contains 6 supplemental Figures and 3 supplemental Tables.

**Supplemental Table 1:** List of genes that are differentially expressed between small intestinal and colonic PDGFRα^hi^ FB ranked in order of significance.

Supplementary Table 2. List of genes that are differentially expressed between small intestinal and colonic Fgfr2^+^ FB ranked in order of significance.

**Supplementary Table 3.** List of common genes that are differentially expressed between both small intestinal and colonic PDGFRα^hi^ FB and small intestinal and colonic Fgfr2^+^ FB.

