## Supplemental information for "The small and large intestine contain transcriptionally related mesenchymal stromal cell subsets that derive from embryonic *Gli1^+^* mesothelial cells"

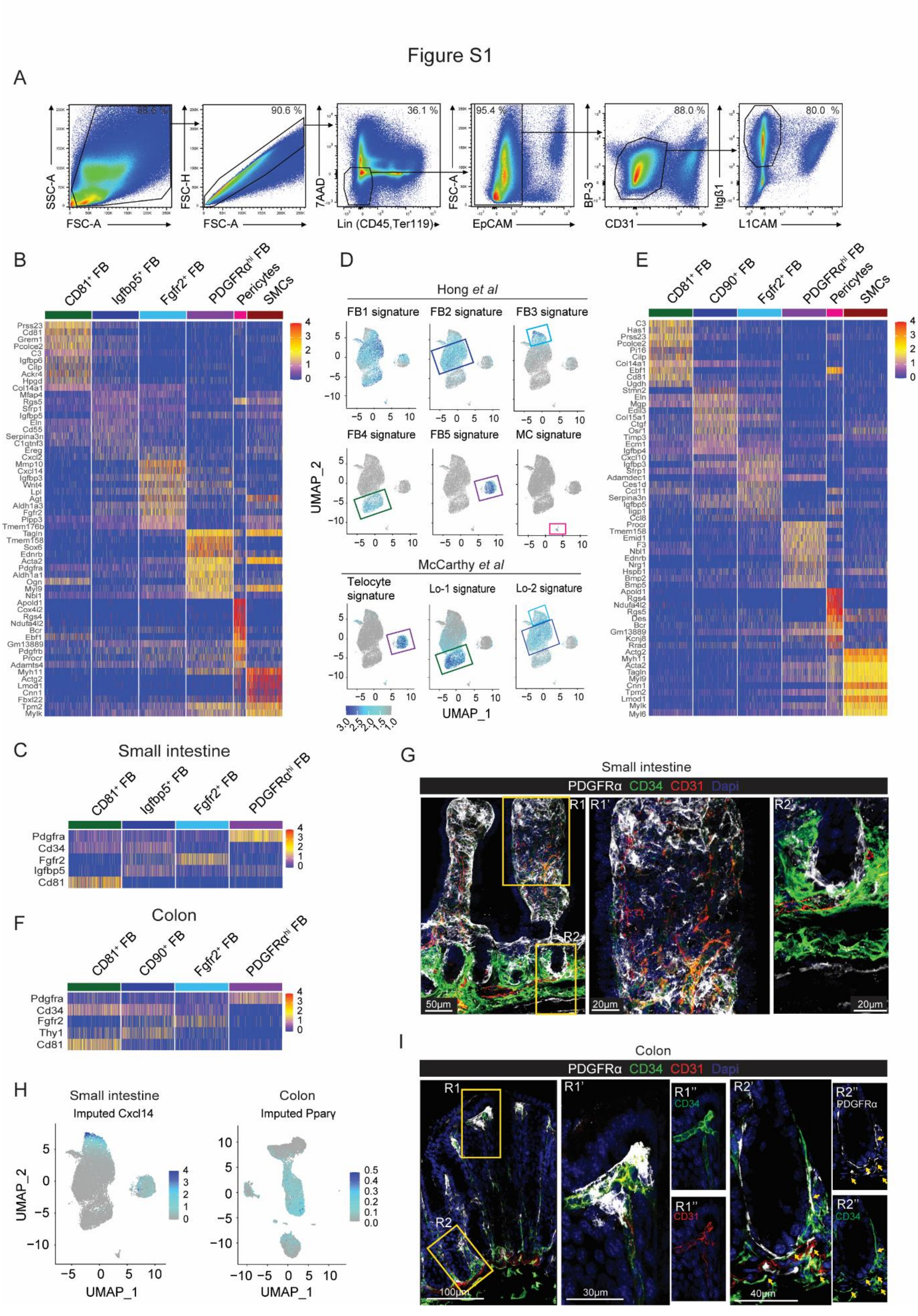

**Figure S1.**

Related to Figure 1. (**A**) Flow cytometric gating strategy for sorting adult small intestinal and colonic Itgβ1$^{+}$ MSC. (**B**) Heatmap with scaled transcription levels (integrated data) of the top 10 differentially expressed genes (DEG) between adult small intestinal MSC subsets. (**C**) Heatmap with scaled transcription levels (anchor integrated data) of markers used to distinguish small intestinal FB subsets. (**D**) Signature genes for MSC subsets identified in Hong *et al* (Hong *et al.*, 2020) and McCarthy *et al* (McCarthy *et al.*, 2020) projected onto the small intestinal MSC UMAP as gene modules. Boxes represent the 6 MSC clusters and are color coded as in (**B**). (**E**) Heatmap with scaled transcription levels (integrated data) of the top 10 DEGs between adult colonic MSC subsets. (**F**) Heatmap with scaled transcription levels (anchor integrated data) of markers used to distinguish colonic FB subsets. (**G)** Immunohistochemical staining of mouse jejunum. R1' and R2' are high magnifications of the R1 and R2 quadrants (yellow squares) in the left image. (**H**) MAGIC imputed projection of (left) *Cxcl14* expression onto the small intestinal MSC UMAP and (right) *Pparγ* expression projected onto the colon MSC UMAP. (**I**) Immunohistochemical staining of mouse colon for indicated antigens. R1'' and R2'' depict single stains. Arrows indicate CD34$^{+}$ FB (CD34$^{+}$PDGFRα$^{+}$CD31$^{-}$ cells). (**H** and **I**) Results are representative stains from (**H**) 3 and (**I**) 2 experiments analyzing intestinal sections from 3 mice/experiment.

Figure S2

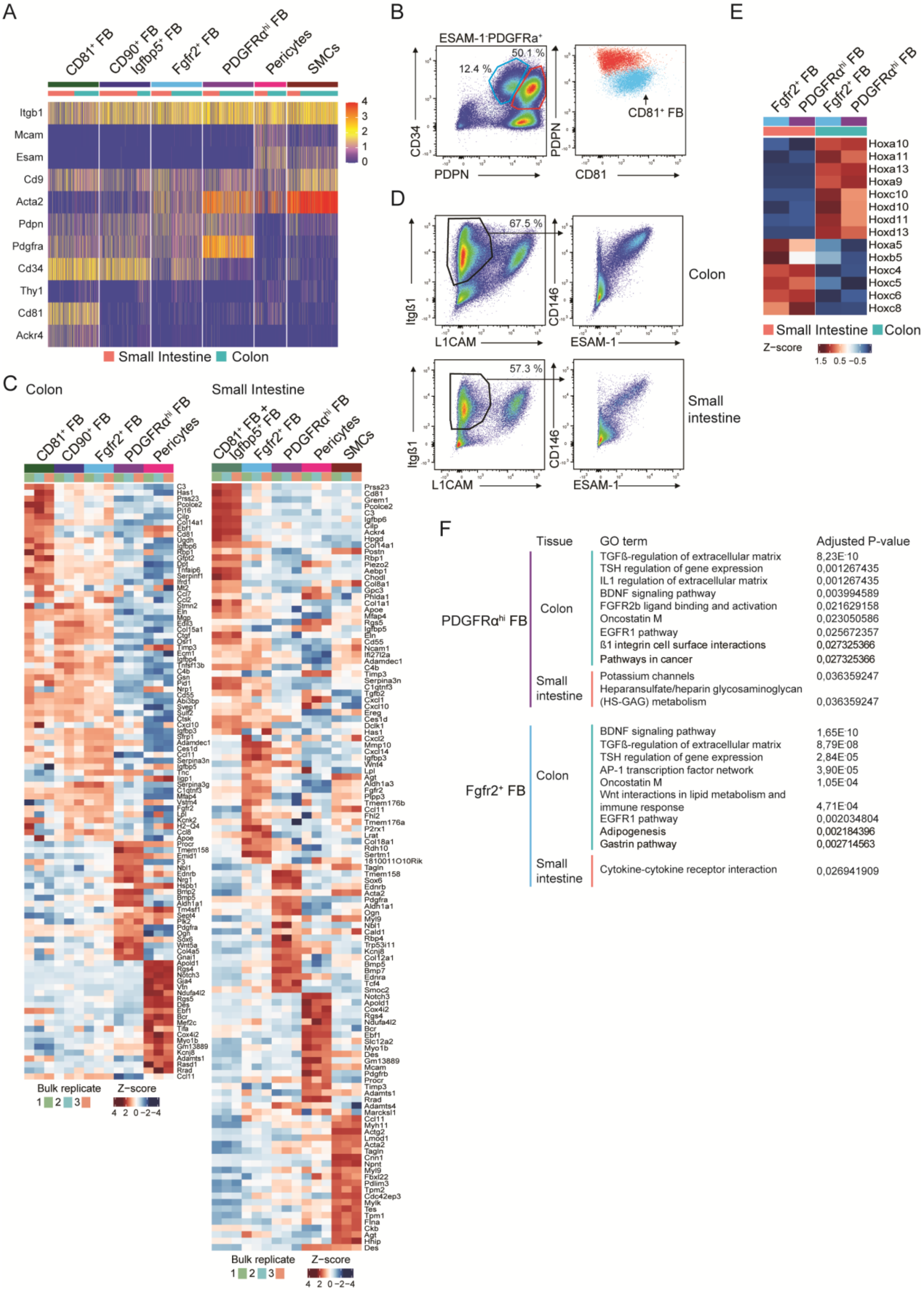
A
CD81+ FB
CD90+ FB
Igfbp5+ FB
Fgfr2+ FB
PDGFRαhi FB
Pericytes
SMCs
Itgb1
Mcam
Esam
Cd9
Acta2
Pdpn
Pdgfra
Cd34
Thy1
Cd81
Ackr4
4
3
2
1
0
Small Intestine
Colon
B
ESAM-1-PDGFRa+
12.4 %
50.1 %
CD34
PDPN
CD81+ FB
CD81
D
67.5 %
Itgß1
L1CAM
CD146
ESAM-1
Colon
57.3 %
Small intestine
E
Fgfr2+ FB
PDGFRαhi FB
Hoxa10
Hoxa11
Hoxa13
Hoxa9
Hoxc10
Hoxd10
Hoxd11
Hoxd13
Hoxa5
Hoxb5
Hoxc4
Hoxc5
Hoxc6
Hoxc8
Small Intestine
Colon
Z-score
1.5 0.5 −0.5
C
Colon
Small Intestine
Bulk replicate
1 2 3
Z-score
4 2 0 −2 −4
F
Tissue
GO term
Adjusted P-value
PDGFRαhi FB
Colon
TGFß-regulation of extracellular matrix 8,23E-10
TSH regulation of gene expression 0,001267435
IL1 regulation of extracellular matrix 0,001267435
BDNF signaling pathway 0,003994589
FGFR2b ligand binding and activation 0,021629158
Oncostatin M 0,023050586
EGFR1 pathway 0,025672357
ß1 integrin cell surface interactions 0,027325366
Pathways in cancer 0,027325366
Small intestine
Potassium channels 0,036359247
Heparansulfate/heparin glycosaminoglycan (HS-GAG) metabolism 0,036359247
Fgfr2+ FB
Colon
BDNF signaling pathway 1,65E-10
TGFß-regulation of extracellular matrix 8,79E-08
TSH regulation of gene expression 2,84E-05
AP-1 transcription factor network 3,90E-05
Oncostatin M 1,05E-04
Wnt interactions in lipid metabolism and immune response 4,71E-04
EGFR1 pathway 0,002034804
Adipogenesis 0,002184396
Gastrin pathway 0,002714563
Small intestine
Cytokine-cytokine receptor interaction 0,026941909

**Figure S2.**

Related to Figure 2. (**A**) Heatmap showing scaled transcription levels (integrated data) of genes used as markers to distinguish the 6 MSC subsets by flow cytometry. (**B**) Flow cytometric analysis verifying that colonic $PDPN^{lo}CD34^{+}$FB (blue gate, left plot) express CD81 (blue cells right plot). Representative gating of at least 3 experiments with 4-7 mice/experiment. (**C**) Heatmaps showing transcription levels of the top 20 DEGs defining the scRNA-seq subsets within the bulk RNA-seq subsets from colon (left) and small intestine (right). (**D**) Flow cytometric gating strategy of colon (top panels) and small intestine (bottom panels) showing co-staining of ESAM-1 and CD146 by $Itg\beta1^{+}$ MSCs. Representative stains of at least 6 experiments with 1 mouse/experiment. (**E**) Heatmap of transcription levels (averaged between bulk RNA-seq triplicates) of Hox genes that were differentially expressed between small and large intestine in the indicated FB subset. (**F**) Gene ontology (GO) analysis using Enrichr (Bioplanet 2019) of DEGs between small intestinal and colonic $PDGFR\alpha^{hi}$ FB (purple) and between small intestinal and colonic $Fgfr2^{+}$ FB (light blue). Top 9 significant pathways based on adjusted p values for colon and all significant pathways for small intestine are shown.

Figure S3

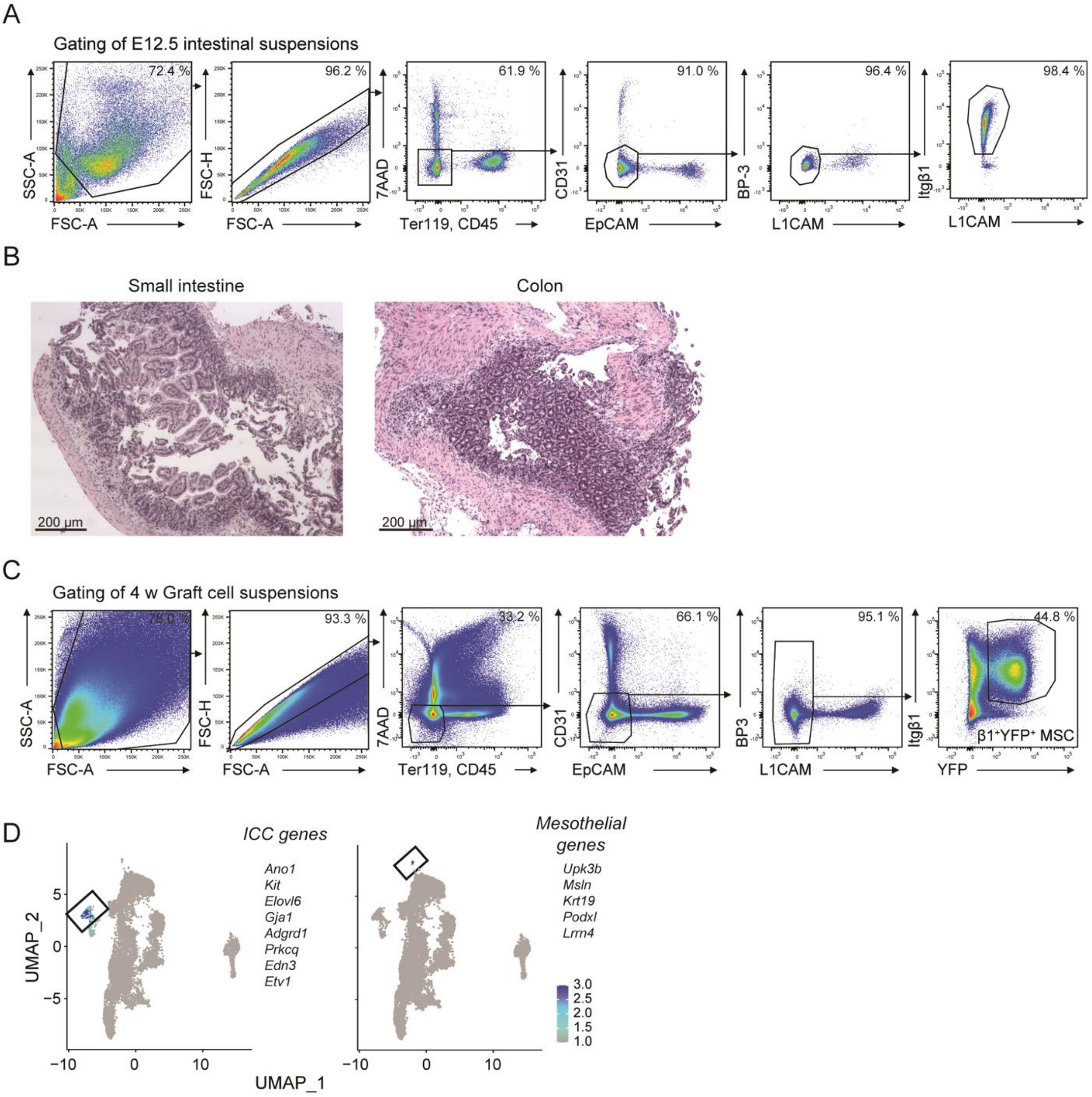


**Figure S3.**

Related to Figure 3. (**A**) Flow cytometric gating strategy for the identification and sorting of intestinal Itgβ1$^+$ MSC from embryonic day (E)12.5 mice. (**B**) Hematoxylin and eosin staining of small intestinal and colonic grafts 6 weeks post transplantation. Results are representative stains of 2 grafts for each tissue. (**C**) Flow cytometric gating strategy for the identification and sorting of YFP$^+$Itgβ1$^+$ MSC from intestinal grafts. (**D**) Projections of module score of signature genes for Interstitial cells of Cajal (ICC) and mesothelium onto colon graft UMAP. Boxes identify the ICC and mesothelial clusters.

Figure S4

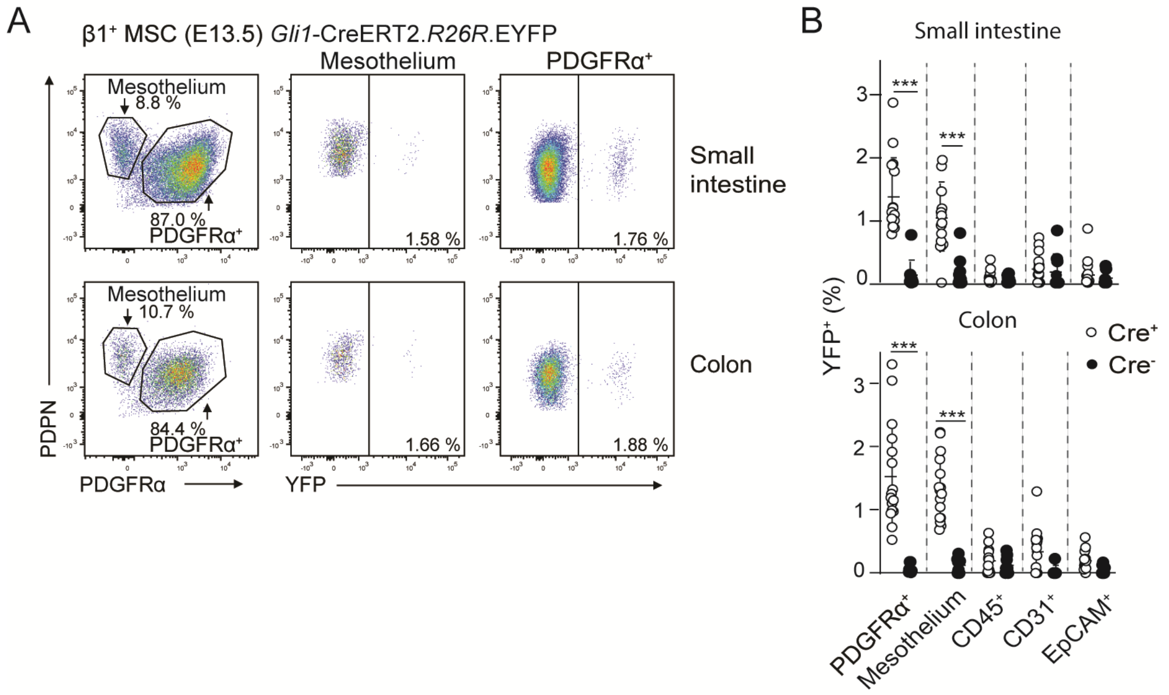


**Figure S4.**

Related to Figure 4. (**A**) Representative flow cytometric analysis and (**B**) pooled data of the proportions of YFP-expressing cells in indicated intestinal populations of E13.5 *Gli1*.CreERT2$^{+/-}$.*R26R*.EYFP mice 2 days after injection with 4-OHT. Pre-gating strategy as in Fig. S3A. Results are from 4 pooled experiments with 1-6 embryos/experiment. Bars, mean (SD). ***p<0.0001, 2-way ANOVA with Benjamini, Krieger and Yekutieli multiple comparisons.

### Figure S5

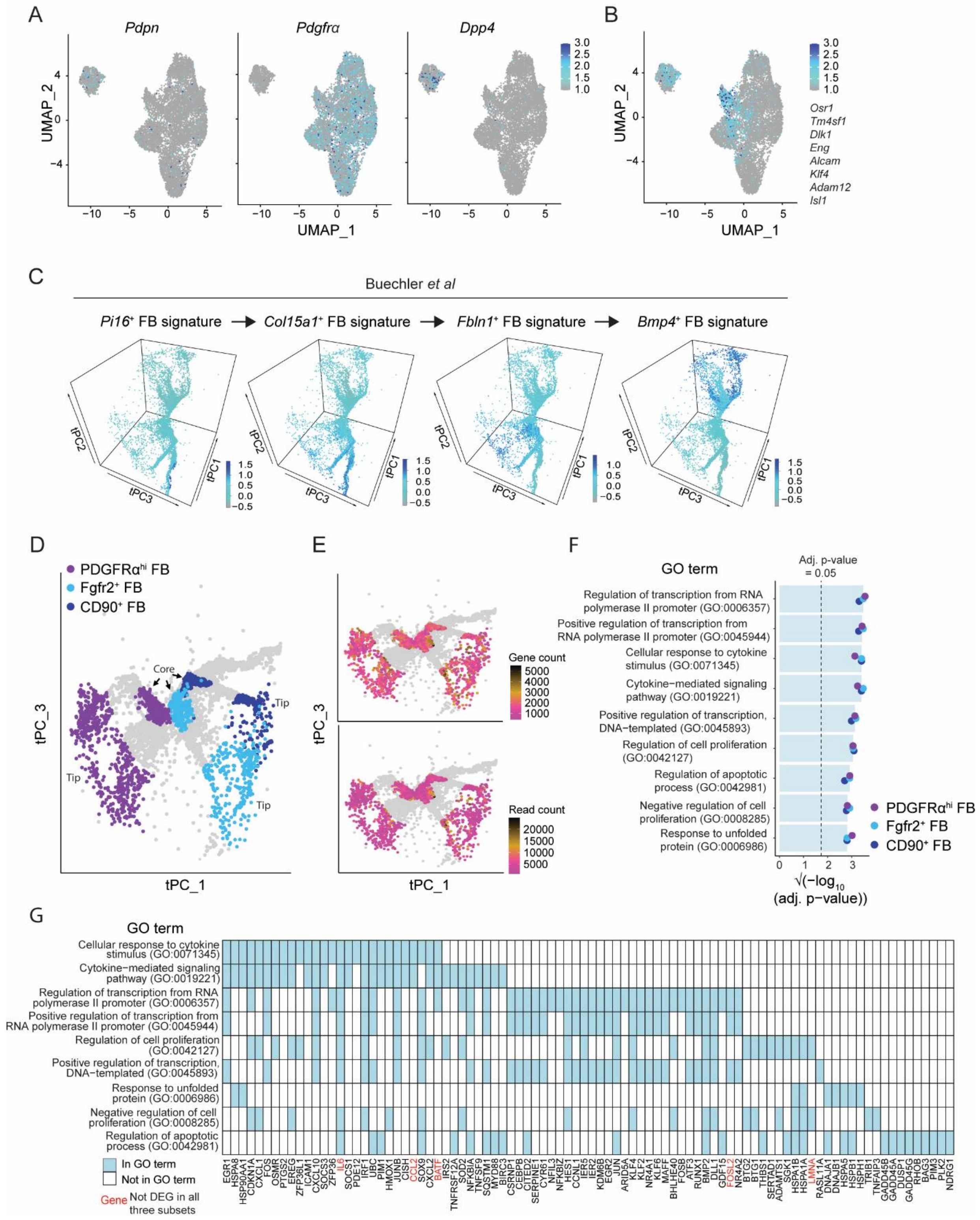

A
Pdpn
Pdgfrα
Dpp4
UMAP_1
UMAP_2
B
Osr1
Tm4sf1
Dlk1
Eng
Alcam
Klf4
Adam12
Isl1
C
Buechler et al
Pi16+ FB signature
Col15a1+ FB signature
Fbln1+ FB signature
Bmp4+ FB signature
D
PDGFRαhi FB
Fgfr2+ FB
CD90+ FB
Core
Tip
tPC_1
tPC_3
E
Gene count
Read count
F
GO term
Adj. p-value = 0.05
Regulation of transcription from RNA polymerase II promoter (GO:0006357)
Positive regulation of transcription from RNA polymerase II promoter (GO:0045944)
Cellular response to cytokine stimulus (GO:0071345)
Cytokine−mediated signaling pathway (GO:0019221)
Positive regulation of transcription, DNA−templated (GO:0045893)
Regulation of cell proliferation (GO:0042127)
Regulation of apoptotic process (GO:0042981)
Negative regulation of cell proliferation (GO:0008285)
Response to unfolded protein (GO:0006986)
√(−log10 (adj. p−value))
G
GO term
In GO term
Not in GO term
Gene Not DEG in all three subsets

**Figure S5.**

Related to Figure 5. (**A**) *Pdpn*, *Dpp4*, *Pdgfrα* integrated and normalized gene expression and (**B**) mesenchymal precursor associated gene expression module score projected onto UMAP of E12.5 large intestinal Itgβ1$^{+}$ MSC. (**C**) Top 20 DEGs for indicated FB clusters identified in Buechler *et al* projected as gene modules onto colonic tSPACE projections of adult colonic MSC in tPC1-3. Arrows indicate predicted trajectory between FB clusters according to Buechler *et al* (Buechler *et al.*, 2021). (**D**) tSPACE principal component analysis projection (tPC1 and tPC3) of adult colonic FB subsets. Colors represent the indicated FB subsets, where core and tip cells within each FB subset were compared for differential gene expression. (**E**) tPC1 vs tPC3 projection showing overlaid gene count (top) and read count (bottom) on tip and core cells highlighted in (**D**). (**F**) GO term analysis using Enrichr (GO Biological Process 2018) of DEGs enriched in core versus tip cells within each FB subset. The data shown are the 9 GO terms shared by core cells within each FB subset (ranked by adjusted p value). Dotted line denotes significant adjusted p value of 0.05. (**G**) Discrete heatmap indicating the DEGs accounting for the top 9 processes. Blue squares represent the genes in GO term. White squares represent the genes not in GO term. Genes labelled black are expressed at significantly higher levels in core compared with tip cells within each FB subsets. Genes in red are expressed at significantly higher levels in core cells from 1-2 of the three FB subsets.

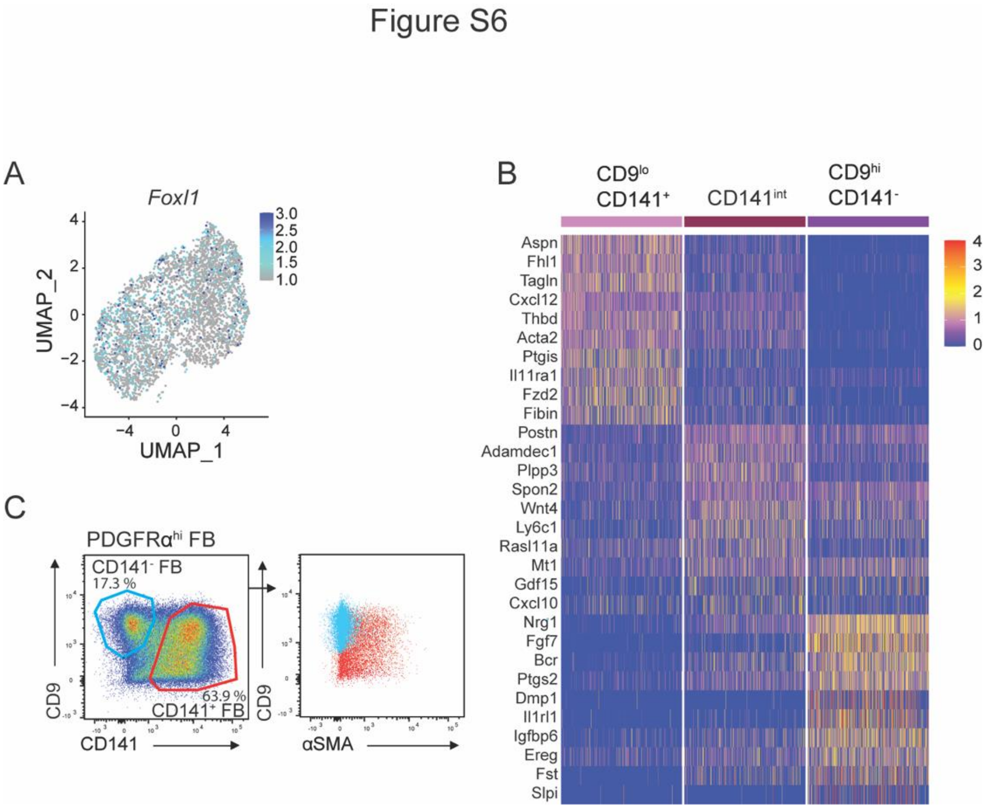


**Figure S6.**

Related to Figure 6. (**A**) Integrated and normalized *Foxl1* expression projected onto UMAP of colonic PDGFRα$^{hi}$ FB. (**B**) Heatmap with scaled expression (integrated) of top 10 differentially expressed genes (DEGs) between the three colonic PDGFRα$^{hi}$ subsets. (**C**) Flow cytometric analysis of PDGFRα$^{hi}$ FB showing expression of αSMA (right hand plot) by CD141$^{+}$ (red) and CD141$^{-}$ (blue) cells using the gates depicted in the left hand panel. Results are representative staining of 2 experiments with 3 mice/experiment.
